# Distinct roles for canonical and variant histone H3 lysine 36 in Polycomb silencing

**DOI:** 10.1101/2022.10.11.511749

**Authors:** Harmony R. Salzler, Vasudha Vandadi, Benjamin D. McMichael, John C. Brown, Sally A. Boerma, Mary P. Leatham-Jensen, Kirsten M. Adams, Michael P. Meers, Jeremy M. Simon, Robert J. Duronio, Daniel J. McKay, A. Gregory Matera

## Abstract

Polycomb complexes regulate cell-type specific gene expression programs through heritable silencing of target genes. Trimethylation of histone H3 lysine 27 (H3K27me3) is essential for this process. Perturbation of H3K36 is thought to interfere with H3K27me3. We show that mutants of *Drosophila* replication-dependent *(H3*.*2*^*K36R*^) or -independent *(H3*.*3*^*K36R*^) histone H3 genes generally maintain Polycomb silencing and reach later stages of development. In contrast, combined *(H3*.*3*^*K36R*^*H3*.*2*^*K36R*^) mutants display widespread Hox gene misexpression and fail to develop past the first larval stage. Chromatin profiling revealed that the *H3*.*2*^*K36R*^ mutation disrupts H3K27me3 levels broadly throughout silenced domains, whereas these regions are mostly unaffected in *H3*.*3*^*K36R*^ animals. Analysis of H3.3 distributions showed that this histone is enriched at presumptive PREs (Polycomb Response Elements) located outside of silenced domains but relatively depleted from those inside. We conclude that H3.2 and H3.3 K36 residues collaborate to repress Hox genes using different mechanisms.

**Short summary:** Histone H3.2 and H3.3 K36 residues ensure Hox gene silencing and enable development by different, but synergistic mechanisms.

## Introduction

A fundamental question in developmental biology is to understand how diverse cell types are generated from undifferentiated precursor cells. Once established, cellular identities must be maintained over time. The failure to do so can result in a wide spectrum of human diseases (*1-4*). Covalent post-translational modifications (PTMs) of the histone proteins that package eukaryotic genomes are thought to encode epigenetic information that is passed from one cell generation to the next (*5-7*), but the mechanisms by which this process occurs remain incompletely understood.

In animal cells, histone PTM functions have largely been inferred from genetic analyses of histone modifying factors (readers, writers, erasers), rather than from studying the histone residues themselves. To help decipher the metazoan ‘histone code’ (*8*), we developed an experimental system in *Drosophila* that allows for sophisticated phenotypic analysis following loss of a specific site of histone modification (*9, 10*). Interestingly, we found that histone missense mutants often exhibit a subset of the phenotypes caused by mutations in their cognate chromatin-modifying enzymes (*9, 11-14*). Here, we take advantage of this system to focus on the role of histone H3 lysine 36 (H3K36) in antagonizing the developmentally regulated gene silencing activity of the Polycomb Repressive Complex 2, PRC2.

A large body of evidence demonstrates that trimethylation of H3 lysine 27 (H3K27me3) is deposited by PRC2 and is critical for formation of silent chromatin (*15, 16*). Recent work has conclusively shown that the H3K27 residue is essential for maintaining repression of homeobox (Hox) genes that control cell fate decisions in *Drosophila* and mice (*9, 17, 18*). Furthermore, allosteric interactions within the PRC2 enzyme complex serve to facilitate spreading of H3K27me3 into neighboring chromatin domains (*19-21*). Thus, the H3K27me3 writer is also a reader – a finding that has profound consequences for understanding the regulation of heterochromatin (*22, 23*). The H3K27me3 mark is also read by Polycomb Repressive Complex 1 (PRC1), containing the Polycomb (Pc) protein, which further condenses and represses H3K27me3 marked genomic regions (*24*).

To counteract the spreading activity of PRC2, other chromatin marks including H3K4me3, H3K36me2 and H3K36me3 are thought to antagonize Polycomb silencing (*25, 26*). Elegant work from Müller and colleagues recently elucidated the structural basis whereby modification of H3K36 inhibits the activity of EZH2, the catalytic subunit of mammalian PRC2 (*27*). Cryo-EM analysis of *in vitro* reconstituted nucleosomes showed that the N-terminal tail of histone H3 is threaded into the active site of EZH2 by a network of interactions that is disrupted by covalent modification of H3K36 (*27*). However, despite the compelling cryo-EM data, histone gene replacement studies *in vivo* in *Drosophila* showed that replication-dependent histone *H3*.*2*^*K36R*^ mutants exhibit reduced H3K27me3 levels but comparatively modest Polycomb derepression phenotypes (*12, 27, 28*). Contrastingly, *H3*.*2*^*K27R*^ mutants exhibit more widespread derepression (*27*). Given the exquisite ability of cells to sense changes in the levels of H3K27-modifiable nucleosomes (*9, 17*), the relatively mild Polycomb phenotypes observed in *H3*.*2*^*K36R*^ larval tissues is puzzling. If the presence of an unmodified H3K36 residue really is necessary for efficient trimethylation of H3K27 *in vivo*, then perhaps there is some redundant histone function that serves to mask the K36R mutant phenotype.

One clear candidate for such a role is the replication-independent histone, H3.3, which differs from H3.2 by only four amino acids. Importantly, the H3.3 and H3.2 N-terminal tails differ by only a single amino acid, at position 31, and thus both can be similarly modified at their respective H3K36 and H3K27 residues (*29, 30*). Accordingly, functional redundancies between H3.2 and H3.3 have been described. One study showed that H3.2 can compensate for loss of H3.3 (*31*). Another example is highlighted by the H3K9 residue (*14*), as the combination of *H3*.*2*^*K9R*^ and *H3*.*3*^*K9R*^ mutations resulted in more severe developmental and transcriptional defects than did either mutation alone.

We therefore hypothesized that lysine 36 of H3.3 might functionally compensate for loss of H3.2K36 with respect to directly promoting E(z) activity, H3K27 trimethylation, and appropriate repression of Hox genes. To test this notion, we generated *H3*.*3*^*K36R*^ mutants and examined them for homeotic phenotypes indicative of faulty Hox gene repression. We also compared the effects of *H3*.*2*^*K36R*^ and *H3*.*3*^*K36R*^ mutations on the levels and genome wide distribution of H3K27me3. These experiments revealed that loss of H3.2K36 causes widespread disruption of H3K27 trimethylation across broad domains (e.g. the Hox gene clusters), whereas loss of H3.3K36 does not. In control genomes, we found that H3.3 preferentially accumulates at presumptive PRC2 recruitment sites, called Polycomb Response Elements, or PREs (*32, 33*). Interestingly, H3.3 accumulates to a lesser degree at PREs located inside broad domains of H3K27me3 silent chromatin than it does to those outside. Finally, we created an *H3*.*2*^*K36R*^*/H3*.*3*^*K36R*^ double mutant and found that combining these mutations synergistically derepresses Hox genes. These findings support a model wherein H3.2K36 and H3.3K36 residues are both important for proper Hox gene repression, but that they carry out this function from distinct genomic subcompartments.

## Results

### Arginine substitutions at K36 and K27 in H3.2 synergistically impair development

The antagonistic relationship between PRC2, which carries out H3K27 trimethylation, and complexes that methylate H3K36 is well established (*25, 26, 34, 35*). However, to date, there is no *direct* evidence demonstrating a developmental biological connection between H3K27 and H3K36 residues. We therefore analyzed animals expressing various bacterial artificial chromosome (BAC) transgenes carrying two homologous copies of a 12x tandemly-arrayed 5kb *Drosophila* histone gene repeat element in the background of a homozygous deletion of the endogenous histone gene cluster (*ΔHisC*), for details see Fig. S1 and Methods. This scheme (Fig. 1A) allowed us to perform genetic complementation analyses by combining 12x transgenes of different genotypes, and thereby assessing the likelihood that different H3.2 residues might participate in a common function. Adults homozygous or hemizygous for the 12x*H3*.*2*^*HWT*^ (histone wild type, *HWT*) tandem array are viable and fertile (*9*), see Figure 1B. We found that one copy of the *HWT* transgene rescues the larval and pupal lethality previously reported (*9, 12*) for 12x*H3*.*2*^*K36R*^ (*K36R*) hemizygotes (Fig. 1B). Interestingly, one copy of the *HWT* transgene was unable to rescue 12x*H3*.*2*^*K27R*^ (*K27R*) hemizygotes. These *K27R/HWT* animals pupate normally, but very few eclose as viable adults (Fig. 1B).

**Figure 1.**
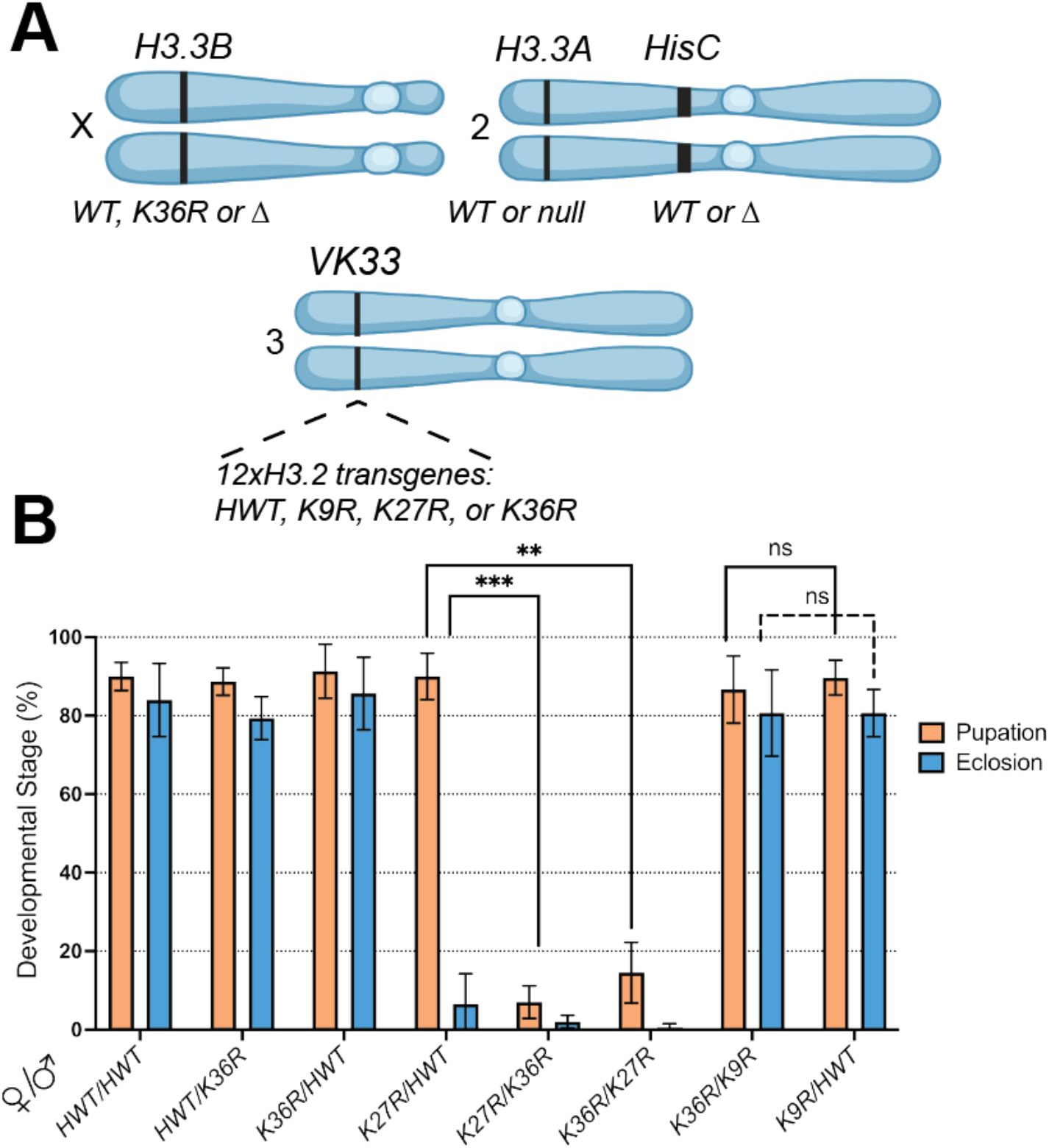
Intragenic complementation analysis within a multi-gene family. **(A)** Cartoon of chromosomal loci used in panel B and in subsequent experiments. For complete genotypes, see Figs. S1, S2 and S4. The *H3.3B* gene (chr. X) is either WT, K36R, or Δ (null). The *H3.3A* gene (chr. 2L) is either WT or Δ (null). The endogenous replication-dependent histone gene cluster *HisC* (chr. 2L) is either intact (WT) or Δ (null). The transgenic insertion site *VK33* (chr. 3L, band 65B2) was used for histone gene replacement analysis. 12xH3.2 transgenes contain 12 copies of each histone repeat unit containing all 5 replication-dependent histone genes. Transgenes used in this study contain the following alleles of *H3.2*: *HWT* (WT), *K9R, K27R*, or *K36R*. Panel created with BioRender.com. **(B)** Developmental viability assay for complementation analysis of *12xH3.2* transgenes. All genotypes are *HisC*Δ and possess two 12x histone transgenes *in trans* (24x total). Pairs of transgenes are represented on the x-axis for each set of bars. For each genotype, %pupation and %eclosion of 4-6 biological replicates (50 larvae/replicate vial) were calculated, and mean and standard deviation (SD) of these percentages were plotted. Statistical significance for %pupation was calculated with GraphPad Prism software using a Mixed-effects analysis (can accommodate missing values) on the 4 genotype pairs indicated by brackets, followed by Šidák’s multiple comparisons test. ** indicates p-value (p) < 0.01; *** indicates p<0.001

Using this assay, we observed a strong genetic interaction between *H3*.*2*^*K36R*^ and *H3*.*2*^*K27R*^; nearly all the *K27R/K36R* animals fail to eclose and most die as larvae before pupation (Fig. 1B). As mentioned above, the control crosses displayed significantly milder phenotypes (Fig. 1B). Notably, the *K27R/K36R* complementation failure is not simply due to an overabundance of mutant histones, as crosses between *K36R* and 12x*H3*.*2*^*K9R*^ (K9R) produced viable adults at similar frequencies to those with *HWT* (Fig. 1B). Previously, this sort of intragenic complementation analysis within a large multi-gene family has not been possible. Our results strongly suggest that H3.2K36 and H3.2K27 residues share common pathways or mechanisms necessary for proper development to adulthood, whereas H3.2K36 and H3.2K9 do not.

### H3.3^K36R^ mutants are viable and fertile

The relative failure of replication-dependent *H3.2*^*K36R*^ mutants to elicit strong Polycomb phenotypes (*12, 27, 28*) suggests the existence of a redundant H3 function. Outside of S-phase and in post-replicative cells, nucleosome turnover is largely carried out using the replication-independent histones, including H3.3 (*36*). In *Drosophila melanogaster*, there are two H3.3 genes, *H3.3A* and *H3.3B* (*37*). To test for genetic redundancies at H3.3 lysine 36, we generated a K to R missense mutation in *H3.3B* using Cas9-mediated homologous recombination, and introgressed it onto an *H3.3A* null mutant (*H3.3A*^*null*^) background (Fig. 1A). Complete genotypes and crossing schemes for generating these animals and all others used in this study can be found in Figs. S1-S4, and Table S1. *H3.3B*^*K36R*^*;H3.3A*^*null*^ double mutant animals (hereafter *H3.3*^*K36R*^) pupate and eclose at similar frequencies to *H3.3A*^*null*^ mutants (Fig. 2A). Thus, the viability of *H3.3B*^*K36R*^ animals was unaffected by deletion of *H3.3A*. Given that animals lacking both *H3.3A* and *H3.3B* (*H3.3Δ*) had previously been shown to complete development (*31, 38*), and those lacking only *H3.3A* are fully viable and fertile (*31*), this result was unsurprising. However, unlike *H3.3*^*K9R*^ or *H3.3*^*K27R*^ mutants (*14, 39*), *H3.3*^*K36R*^ males are fertile. Furthermore, these data reveal that the H3.3K36R protein is incorporated into chromatin and is at least partially functional, as *H3.3*^*K36R*^ mutants eclose at significantly higher frequencies than *H3.3Δ* animals (Fig. 2A).

**Figure 2.**
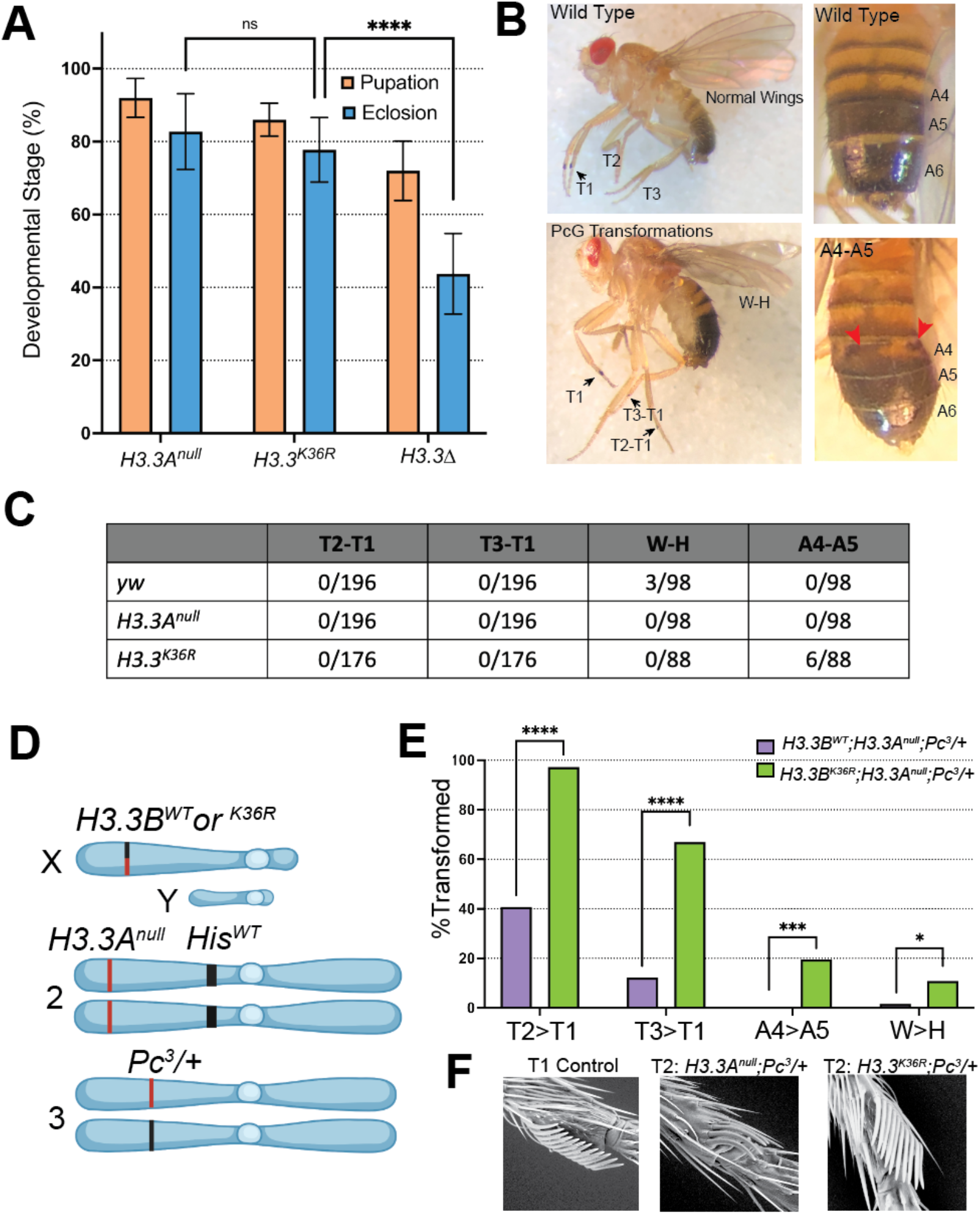
Homeotic transformation analysis of *H3.3*^*K36R*^ mutants. **(A)** Developmental viability of *H3.3*^*K36R*^ mutant and controls. *H3.3*^*K36R*^ indicates *H3.3A*^*null*^ combined with *H3.3B*^*K36R*^. Assay conditions and statistical analyses are as in Figure 1B, with the following modifications. 8 biological replicates were scored for all genotypes. Statistical significance was calculated on %eclosion with One-way ANOVA, as there were no missing values. Šidák’s multiple comparisons test was used as above, with the following additional p-values. * = p<0.05; ****=p<0.0001. **(B)** Top left panel: a wild type control fly with normal legs and wings. Bottom left panel: a fly with 3 homeotic transformations scored in panel C: T2-T1 (thoracic segment leg 2 to leg 1), T3-T1 (leg 3 to leg 1), and W-H (wing to haltere). Top right panel: a fly with a wild type abdomen. Bottom right panel: a fly exhibiting a typical A4-A5 (abdominal segment 4 to abdominal segment 5) transformation scored in this assay. Individual A4-A6 segments are labeled. Red arrows highlight areas of abnormal pigmentation indicating a partial A4-A5 transformation. **(C)** Table showing number of transformations per scored events for each *H3.3*^*K36R*^ mutant and control genotype. **(D)** *H3.3*^*K36R*^ (and control genotypes) were combined with a heterozygous Pc^3^ mutation and scored for 4 characteristic PcG homeotic transformations depicted above in B. For full genetic scheme, see Fig. S3. This panel was created using BioRender.com. **(E)** % Transformed for these transformations is plotted for each genotype. N value for number of flies scored for the *H3.2*^*K36R*^ genotype (n=55) and for the control (n=62). Note, for T2-T1 and T3-T1, each appendage was scored separately, effectively doubling the n value for these transformations. GraphPad Prism was used to calculate a χ^2^ value for each transformation. Significance is abbreviated as: *=p<0.05, **=p<0.01, ***=p<0.001, ****=p<0.0001. **(F)** Image of a typical T2-T1 transformation for each genotype collected by scanning electron microscopy at 250x magnification.

### The H3.3K36R mutation enhances Polycomb phenotypes in adults

The *H3.3*^*WT*^*H3.2*^*K36R*^ mutants exhibit delayed development, with a broad lethal phase that extends throughout larval and pupal stages; very rarely (< 0.2%) animals eclose as adults ((*9, 12*); our unpublished observations). However, *H3.3*^*WT*^*H3.2*^*K36R*^ pharate adult (uneclosed) males demonstrate homeotic transformations that are indicative of impaired repression of Hox genes (*27*). These include: thoracic T2 to T1 (T2-T1), and T3 to T1 (T3-T1), abdominal segment 4 to segment 5 (A4-A5), and wing to haltere (W-H) transformations. Phenotypically, the T2-T1 and T3-T1 leg transformations display ectopic sex comb bristles (Fig. 2B), the A4-A5 transformations show abnormal pigmentation of segment A4 (Fig. 2B), and the W-H transformations feature abnormal wing morphology, manifesting as fully or partially crumpled wings (Fig. 2).

Control *yw* flies (n=98) with wild type histone loci completely lacked leg and abdominal transformations, though 3/98 flies displayed wing morphology consistent with mild W-H transformations. Inspection of *H3.3*^*K36R*^ adult males (n=88) for homeotic transformations revealed few overt PcG phenotypes compared to controls (Fig. 2C, (*40, 41*)). However, introducing *H3.3*^*K36R*^ into a sensitized genetic background revealed a significant increase in homeotic transformation. Flies heterozygous for a null mutation of *Pc* (*Pc*^*3*^*/+*), are viable as adults, but exhibit a baseline frequency of homeotic transformations, which can be modified by other mutations present in the genotype (*42-44*). This assay has been used reliably to implicate genes regulating Pc dependent gene silencing in many other studies (*42-44*). We constructed two pairs of control and mutant genotypes for analysis (Fig. 2D-F). As the full genotypes are quite lengthy, we have abbreviated them for clarity. See Fig. S3 for complete genotypes.

First, to validate the role of H3.2K36 in Hox gene repression in this assay, and to provide a benchmark for subsequent analysis of H3.3, we scored T2-T1, T3-T1, W-H, and A4-A5 transformations in *HisΔ/+; H3.2*^*K36R*^*/Pc*^*3*^ mutants and *HisΔ/+; H3.2*^*HWT*^*/Pc*^*3*^ controls (Fig. S5). Despite mutant *H3.2*^*K36R*^ genes comprising only about 10% of the total number of *H3.2* genes (*9, 45*), we observed significant increases in L2-L1, L3-L1, and W-H transformations in the *HisΔ/+; H3.2*^*K36R*^*/Pc*^*3*^ mutants compared to *HisΔ/+; H3.2*^*HWT*^*/Pc*^*3*^ control animals (Fig. S5). These data validate this assay in the context of histone N-terminal tail residue mutation, provide further evidence that the H3.2K36 residue promotes Pc mediated silencing of Hox genes, and provide a benchmark for the sensitivity of this assay.

Second, we further investigated whether H3.3K36 plays a role in Pc mediated Hox gene repression by scoring homeotic transformations in *H3.3*^*K36R*^*;Pc*^*3*^*/+* mutants and *H3.3A*^*null*^ *;Pc*^*3*^*/+* controls (Fig. 2D-E). Of note, these flies had a fully wild type *HisC* locus, and thus all replication-dependent *H3.2* genes were present at wild-type copy number. We observed a significant increase in all four categories of homeotic transformations in the *H3.3*^*K36R*^*;Pc*^*3*^*/+* mutants relative to the control group (Fig. 2E). Interestingly, we observed a sizeable increase in A4-A5 transformations (19.6%, p<0.001) when *H3.3* was mutated, compared to when *H3.2* was mutated (3.6%, p=ns). This observation implies that the A4-A5 transformation is particularly sensitive to H3.3K36 mutation. Such relative differences in the severity of homeotic transformation between *H3.2*^*K36R*^ and *H3.3*^*K36R*^ mutants suggest that H3.2K36 and H3.3K36 might promote Hox gene repression by non-identical means.

In the above assay, we quantitatively scored homeotic transformations as differences in frequency. However, we also noted qualitative differences in phenotypic severity that mirrored or exceeded these changes in frequency. For example, while *H3.3B*^*K36R*^; *H3.3A*^*null*^*;Pc*^*3*^*/+* mutant and *H3.3B*^*WT*^*;H3.3A*^*null*^*;Pc*^*3*^*/+* controls both exhibit T2-T1 sex comb transformations, the *H3.3B*^*K36R*^; *H3.3A*^*null*^*;Pc*^*3*^*/+* mutant generally demonstrated a more severe phenotype, as indicated by the number of bristles on T2. To capture these qualitative differences in sex comb transformations, we utilized scanning electron microscopy to image T2 sex combs of each genotype in the previous sets of experiments (Fig. 2F). We consistently observed a greater number of bristles in the T2 sex combs of both *HisΔ/+; H3.2*^*K36R*^*/Pc*^*3*^ and *H3.3*^*K36R*^*;Pc*^*3*^*/+* mutant genotypes relative to controls (Fig. 2F). These same relative differences in severity also applied to the other transformations that we scored quantitatively. In summary, these analyses of adult animals suggest roles for both H3.2K36 and H3.3K36 in repression of Hox genes at late developmental stages.

### H3.3K36 and H3.2K36 differentially affect H3K27me3 late in development

Given that PcG phenotypes arise in *H3.3*^*K36R*^ adults, we wondered whether an effect on H3K27me3 might become evident at this later developmental time point. Therefore, we next sought to measure the relative impact of H3.2 and H3.3 K36R on H3K27me3 by western blotting from extracts of adult heads. If unmethylated H3.3K36 were redundantly stimulating E(z) at later developmental stages, we would expect to observe reduced H3K27me3 in *H3.3*^*K36R*^*;Pc*^*3*^*/+* mutants relative to *H3.3A*^*null*^ *;Pc*^*3*^*/+* controls, and relative to the *HisΔ/+;H3.2*^*K36R*^*/Pc*^*3*^ genotype. However, we did not observe any difference in global H3K27me3 levels in the *H3.3*^*K36R*^*;Pc*^*3*^*/+* animals compared to control, despite a clear effect on homeotic transformation frequency (Figs. 3, 2E). In contrast, we observed a ∼40% decrease in H3K27me3 in *HisΔ/+; H3.2*^*K36R*^*/Pc*^*3*^ mutants relative to *HisΔ/+; H3.2*^*HWT*^*/Pc*^*3*^ controls, despite the fact that only 10% of *H3.2* histone genes carry the *H3.2*^*K36R*^ mutation in this gentoype (Figs. 3, S5). This large difference held true in 3 separate experiments (p<0.05, Fig. 3B). Overall, these data are consistent with the possibility that H3.2K36 is more important than H3.3K36 in promoting global levels of H3K27me3 even at a late developmental timepoint.

**Figure 3.**
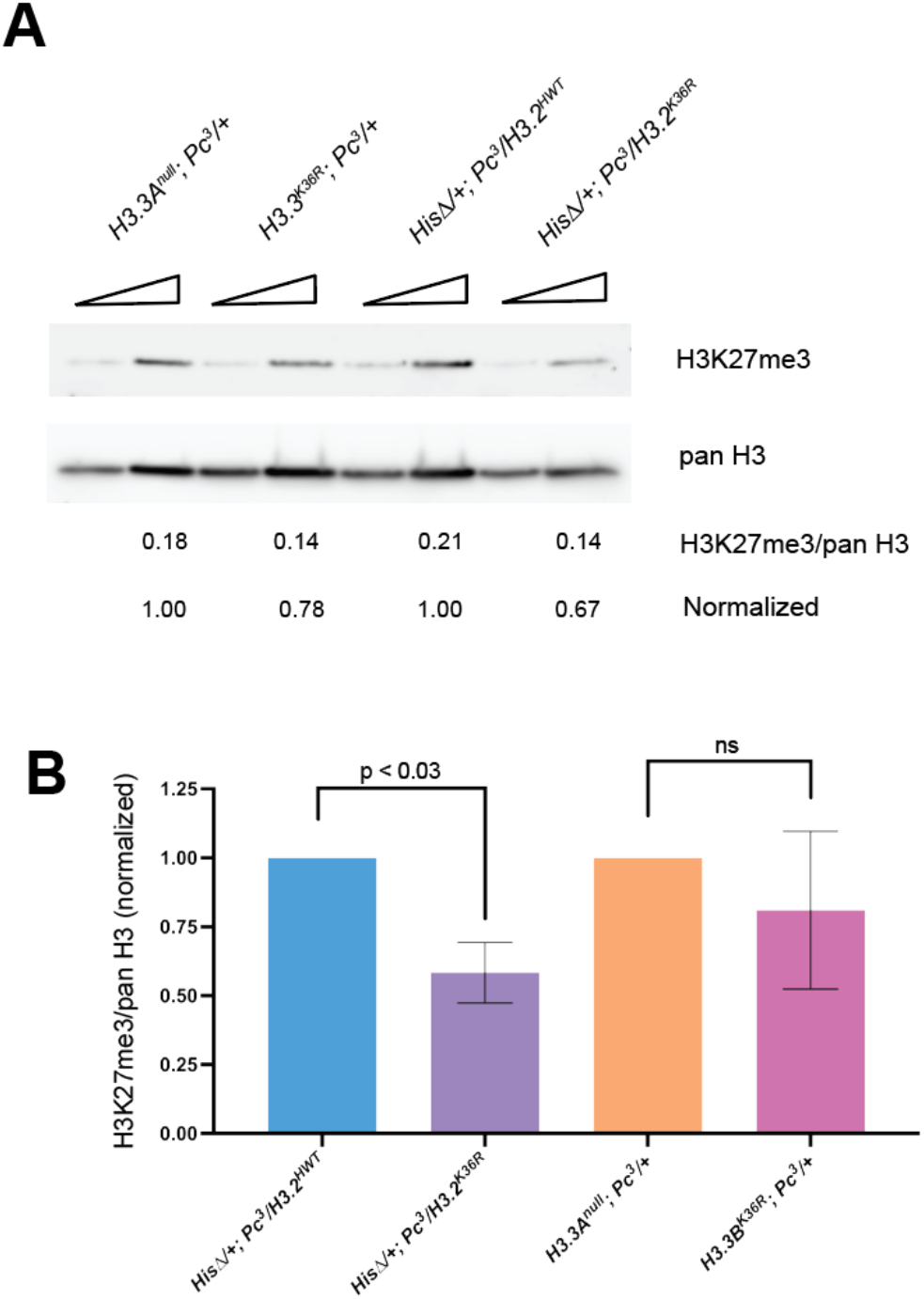
Western blot analysis of H3K27me3 levels in adult heads. *H3.2* and *H3.3* K36R mutants were maintaned in a sensitized *Pc*^*3*^ genetic background. **(A)** Western blotting with anti-H3K27me3 and -panH3 were performed using adult head extracts, loading either 40 or 80ug of protein per lane. Three independent experiments were performed and imaged; a representative blot is shown. Mean pixel intensity was calculated for each band using equal sized areas, and a ratio of H3K27me3/panH3 was calculated for each biological replicate from the 80ug lane. **(B)** For each independent experiment, the H3K27me3/panH3 ratio of mutant genotypes was normalized to each control (*ΔHis/+; H3.2* ^*K36R*^ */Pc*^*3*^ to *ΔHis/+; H3.2* ^*HWT*^ */Pc*^*3*^; *H3.3* ^*K36R*^; *Pc*^*3*^*/+* to *H3.3A*^*null*^; *Pc*^*3*^*/+*). For each genotype, the normalized mean and SD of H3K27me3/panH3 values of all 3 experiments was plotted. Statistical significance between raw H3K27me3/H3 ratios was assessed by paired one way ANOVA (within experiments), followed by Friedman Tests individually comparing mutant genotypes to controls. P-values are directly noted on the graph.

### Mutation of H3.2K36, but not H3.3K36, causes defects in H3K27me3 spreading

Previous studies in various organisms have revealed antagonism between factors that carry out H3K36me and H3K27me. In particular, H3K36me is thought to be important for demarcating Pc domain boundaries (*34, 46-48*). To investigate changes in H3K27me3 patterns in the *H3.2*^*K36R*^ and *H3.3*^*K36R*^ mutants, we employed CUT&RUN chromatin profiling in wing imaginal discs of wandering 3^rd^ instar (WL3) larvae (*49*). To directly compare H3K27me3 levels between genotypes, we analyzed the *H3.3*^*K36R*^ mutation in the same genetic background used for the histone gene replacement platform. The resultant *H3.3*^*K36R*^*H3.2*^*HWT*^ animals were compared with *H3.3A*^*null*^*H3.2*^*HWT*^ controls, each of which contain a deletion of the *H3.3A* gene (see Fig. S4 for details). For each mutant and control, we performed 3 independent biological replicates, and sequenced both the supernatant and pellet fractions (see Methods). The data in both fractions were consistent with respect to signal distribution, however correlations between samples of the same genotype were superior in the pellet fraction (Fig. S7), Thus for all subsequent analyses with the anti-H3K27me3 antibody, we utilized the pellet fraction, except where specifically noted.

As expected, H3K27me3 was highly enriched across known Pc domains in all genotypes, exemplified by the browser shot of the Bithorax complex (Fig. 4A). We noted no discernable difference between the two control genotypes. Genome-wide, we also observed little difference between the *H3.3*^*K36R*^*H3.2*^*HWT*^ mutant and the *H3.3A*^*null*^*H3.2*^*HWT*^ control (Fig. 4A). In contrast, H3K27me3 levels in the *H3.3*^*WT*^*H3.2*^*K36R*^ mutant were markedly depleted relative to the *H3.3*^*WT*^*H3.2* ^*HWT*^ control (Fig. 4A). To quantify the number of broad H3K27me3 domains that changed for each mutant, we performed a differential peak analysis using DESeq2 (Fig. 4B). These domains spanned a range of sizes and H3K27me3 signal intensities, from very large and heavily methylated regions such as the Bithorax and Antennapedia complexes, to much smaller and less intensely methylated ones (Fig. 4B). In the *H3.3*^*K36R*^*H3.2*^*HWT*^ mutants, there were very few (9/677) differential domains with an adjusted p-value <0.05 and a Log_2_ fold-change (LFC) threshold of > |1| (Fig. 4B). Moreover, the nine differential peaks identified in the *H3.3*^*K36R*^*H3.2*^*HWT*^ animals all displayed *increased* H3K27me3 signal intensity in the mutant, which is the opposite of what would occur if K36R were inhibiting PRC2. In contrast, the *H3.3*^*WT*^*H3.2*^*K36R*^ mutants exhibited significantly more differentially methylated domains (169/677), the majority of which (116/169) decreased in intensity (Fig. 4B) (*27*). Indeed, the most striking changes are evident in domains bearing the highest levels of methylation (Base Mean >10^3^), where nearly all of them decreased in the presence of the mutation (Fig. 4B). Overall, these data provide no evidence that H3.3K36 contributes to methylation within broad H3K27me3 domains in the wing disc, whereas H3.2K36 has a widespread effect.

**Figure 4.**
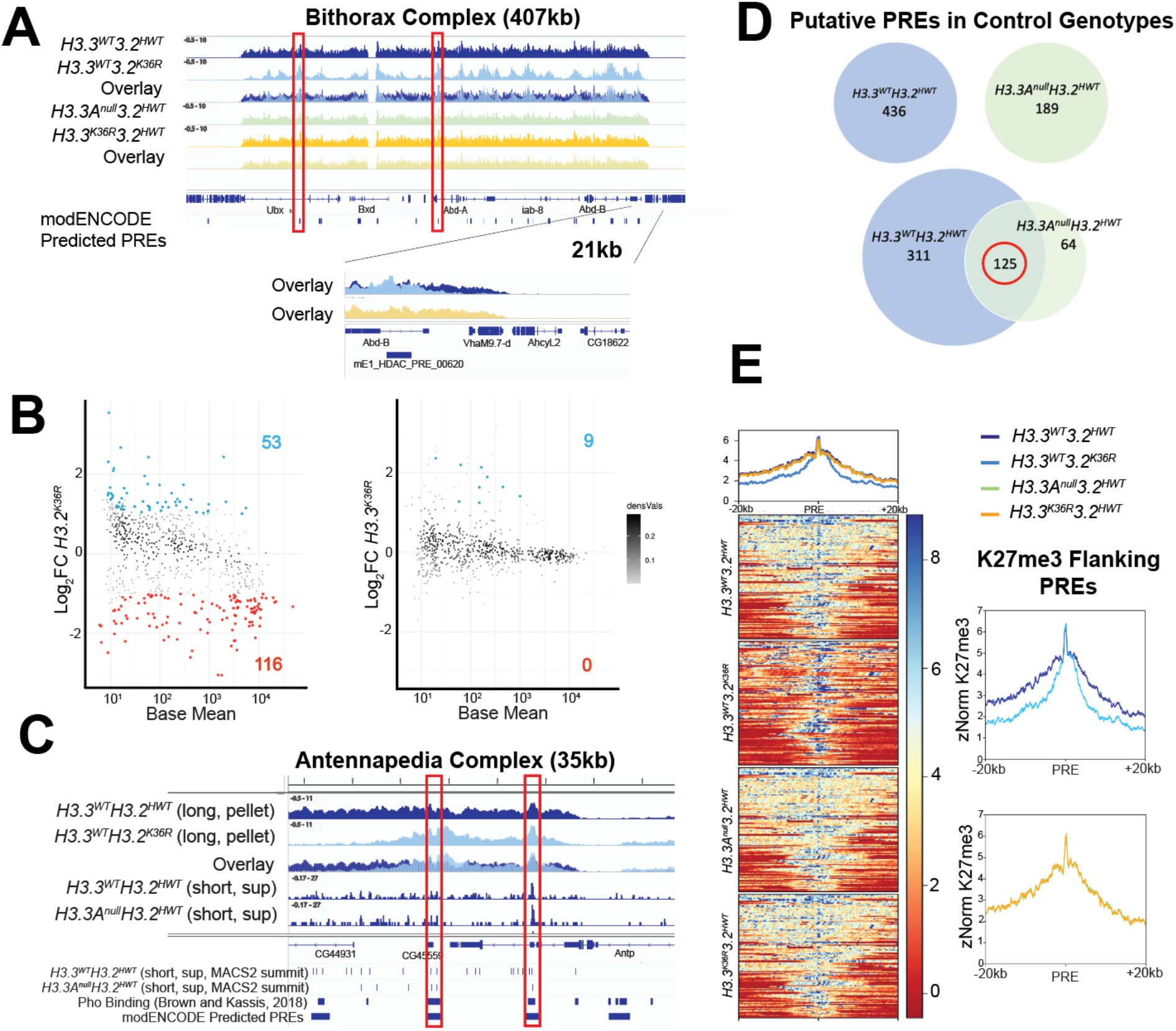
Mutation of H3.2K36, but not H3.3K36, impairs H3K27me3 in Polycomb domains. CUT&RUN profiling of H3K27me3 was performed in WL3 wing disc tissue from three biological replicates of mutant and control genotypes. All browser tracks, metaplots, and heatmaps utilized z-score normalization **(A)** An IGV browser shot of the Bithorax Complex from pellet fraction merged bigwig files. Individual and overlay tracks for *H3.3*^*WT*^*H3.3*^*K36R*^ and *H3.3*^*WT*^*H3.3*^*HWT*^ control, and *H3.3*^*K36R*^*H3.3*^*HWT*^ and *H3.3A*^*null*^*H3.3*^*HWT*^ control are represented. Below the genome annotation track are PREs predicted by modENCODE. Red boxes indicate predicted PREs where all genotypes have wild type H3K27me3, with H3K27me3 declining in the *H3.3*^*WT*^*H3.3*^*K36R*^ mutant in flanking regions. A 21kb region containing the AbdB promoter is shown in higher magnification at the bottom. **(B)** Differential peak analysis of broad H3K27me3 domains using DESeq2. For this comparison, all MACS2 broad peak intervals within 10kb of one another were further merged prior to running DESeq2 (see Fig. S8). Points with an adjusted p-value > 0.05 and a Log_2_ fold change (Log_2_FC) > |1| are shaded (red = downregulated, blue = upregulated). Log_2_FC is computed relative to the appropriate control based on its *H3.3A* status. **(C)** An IGV browser shot using merged bigwig files of H3K27me3 from the Antennapedia Complex using reads from long length fragments (150-700bp). Tracks are shown for the *H3.3*^*WT*^*H3.3*^*K36R*^ mutant and *H3.3*^*WT*^*H3.3*^*HWT*^ control, including an overlay. Below the overlay, tracks from sub-nucleosomal short (20-120bp) fragments of the CUT&RUN supernatant fraction from the two control genotypes are shown. BED tracks at the bottom represent individual MACS2 narrow peak summits, Pho binding sites as determined in Ref. (*59*), and modENCODE predicted PREs. Red boxes indicate correspondence between these features. **(D)** To determine putative PREs, intervals containing MACS2 peak summits + 150bp that overlapped with Pho binding regions (see Methods and Fig. S8) were identified for each control. The number of intervals for each control genotype is indicated in each circle (*H3.3*^*WT*^*H3.3*^*HWT*^, blue; *H3.3A*^*null*^*H3.3*^*HWT*^, green). We utilized the intersection of PRE intervals identified in both controls (circled in red) for further analysis. **(E)** Heatmap and metaplots of H3K27me3 signal intensity at putative PREs and 20kb flanking regions determined in panel D, calculated from merged bigwig files from each genotype. To the right, each mutant genotype is compared directly to its respective control.

The pattern of depletion in the *H3.3*^*WT*^*H3.2*^*K36R*^ mutant was also striking. That is, the ‘spikes’ of K27 trimethylation in the mutant were of comparable magnitude to those in the controls, but the signal was severely depleted in the troughs, and at Pc domain boundaries (Fig. 4A). One might expect a spiky pattern such as this if PRC2 could efficiently nucleate at specific sites (e.g., PREs) but was unable to spread effectively between them. Although PREs are generally thought to be nucleosome depleted (*32, 50*), DNA binding factors located at these sites are known to recruit PRC2 to adjacent nucleosomes. During the CUT&RUN assay, DNA cleavage by micrococcal nuclease (MNase) is directed by an antibody. In this case, anti-H3K27me3 is expected to recruit MNase to cleave chromatin that is near PREs into short, sub-nucleosomal fragments (Fig. 5A, (*50-52*)). Accordingly, the spikes located within broader domains of H3K27me3 signal in our data align well with PREs that have been predicted by other methods (Fig. 4A, (*53*)).

**Figure 5.**
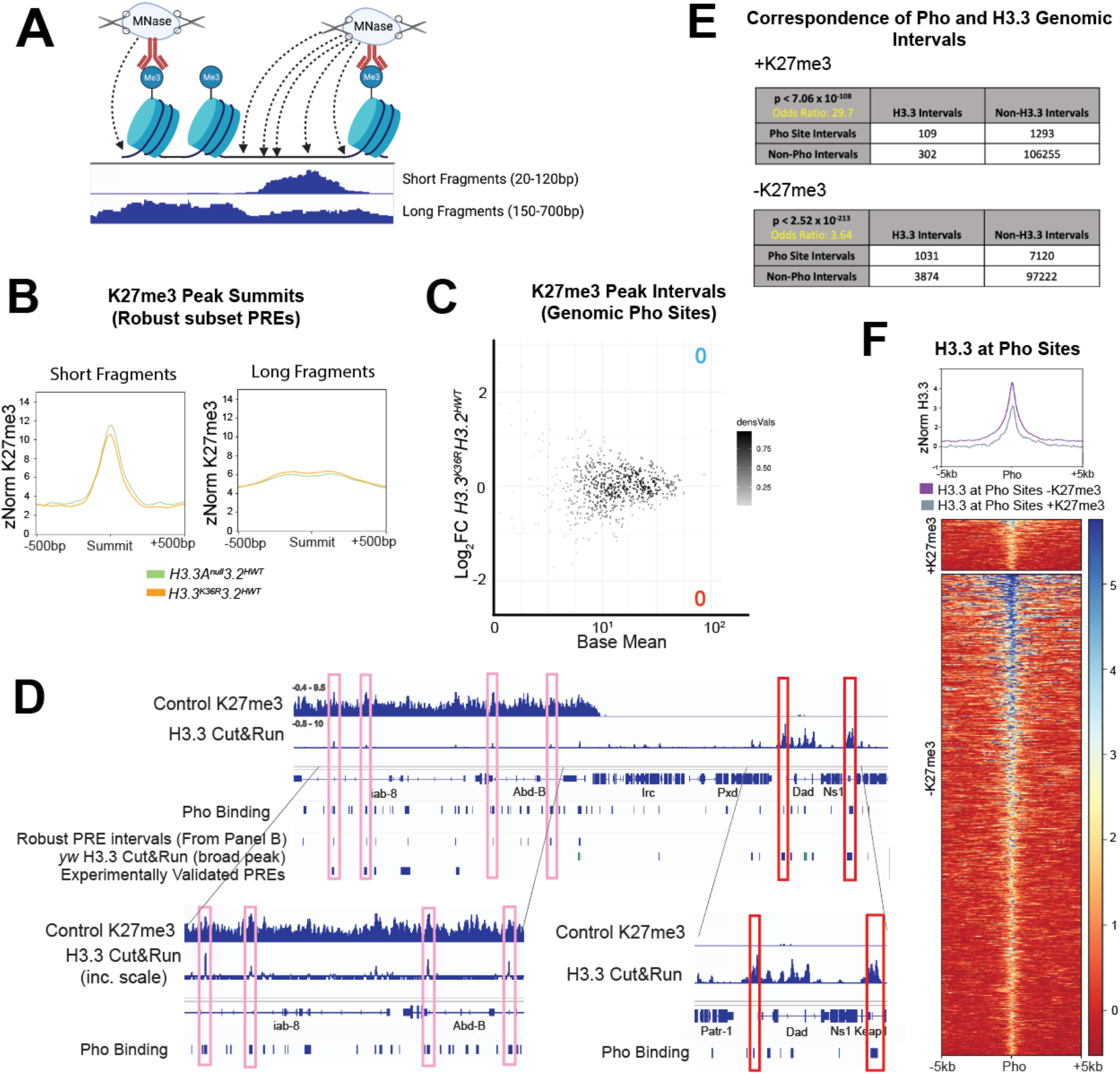
H3K27me3 directed cleavage at presumptive PREs is unchanged in *H3.3*^*K36R*^ mutants. **(A)** Cartoon of MNase cleavage near histone PTMs during CUT&RUN. Antibodies direct the localization of MNase, which cleaves nearby accessible DNA to generate both long (nucleosomal) and short (sub-nucleosomal) fragments. Image produced using Biorender.com **(B)** Metaplots of K27me3 directed CUT&RUN signal + 500bp around peak summits, called from all four genotypes, that overlap with the robust PRE intervals identified in Fig. 4D (n=344). Separate plots were generated for long and short fragments. The *H3.3*^*K36R*^*H3.2*^*HWT*^ mutant is shown alongside its *H3.3A*^*null*^*H3.2*^*HWT*^ control. **(C)** DESeq2 differential analysis of H3K27me3 peak summit intervals that overlap with genome-wide Pho binding sites identified by Kassis and colleagues (n=711). An M/A plot of these intervals for the *H3.3*^*K36R*^*H3.2*^*HWT*^ mutant vs. *H3.3A*^*null*^*H3.2*^*HWT*^ control is shown (details in Fig. S8). No significant peaks were identified. **(D)** An IGV browser shot of H3.3 CUT&RUN at the Bithorax Complex from supernatant fraction, merged, z-score normalized bigwig files. Two replicates each of a *yw* control bearing no histone mutations were performed to determine H3.3 signal, and an *H3.3Δ* mutant was used to subtract regions with non-specific antibody binding. An H3K27me3 track is included for identification of H3K27me3 domains. BED tracks are included below that show positions of: Pho binding sites identified in Ref. (*59*); robust PRE intervals derived in panel B; MACS2 peak calls from H3.3 CUT&RUN from *yw* control genotype; experimentally validated PREs curated in Ref. (*54*). Lighter (pink) boxes illustrate subthreshold H3.3 peaks within the Bithorax complex. Darker (red) boxes show H3.3 peaks in the vicinity of the nearby *Dad* locus. **(E)** Fisher’s Exact Tests measuring correspondence between genomic intervals encompassing Pho sites and those encompassing H3.3 binding. Separate tests were performed for regions inside and outside H3K27me3 domains. Correspondence was highly significant in both cases. **(F)** Metaplot and heatmap comparing z-normalized H3.3 CUT&RUN signal in *yw* controls at Pho sites located inside (+K27me3) and outside (-K27me3) of H3K27me3 domains.

The depletion of H3K27me3 signal between presumptive PREs in the Hox gene clusters suggests that the *H3.3*^*WT*^*H3.2*^*K36R*^ mutation causes a defect in spreading of Pc silencing factors. PREs are tissue-specific cis-regulatory modules, so to determine if the spreading defect is a general phenomenon, we first needed to identify a robust set of PREs that are utilized in the WL3 wing disc. The gold standard definition of a PRE is a functional one, requiring experimental characterization of individual sequences *in vivo*. Thus, a relatively small number of PREs have been validated in this manner, and most of the experiments have not been performed in wing discs (*32, 54*). Bioinformatic predictions based on the presence of DNA binding protein motifs (*55-58*) predict elements with potential PRE activity but do not define which sequences are utilized in a particular developmental scenario. To circumvent these problems, we set out to identify PREs utilized in the WL3 wing disc in a manner that is both functional and predictive.

As functional PREs encompass sequences where DNA binding proteins assemble at open regions of chromatin, we exploited the propensity for accessible chromatin to be cleaved into sub-nucleosomal fragments in the CUT&RUN assay (Fig. 5A, (*52*)) by centering our analysis on peaks derived from short fragments (20-120bp) located within broader H3K27me3 domains (Fig. 4C). For this purpose, we utilized MACS2 narrow peak summit calls from short fragments in the supernatant fraction, as the supernatant yielded fewer spurious peak calls due to localized noise. To improve accuracy, we also capitalized on the availability of ChIP-seq data that were generated from a mixed sample of larval brain and imaginal discs using antibodies targeting pleiohomeotic (Pho; (*59*)). In *Drosophila*, the presence of Pho is one of the best predictors of functional PRC activity at experimentally verified PREs (*32, 60*). Using these two data sets, we bioinformatically identified genomic intervals of short-fragment H3K27me3 peak summits (+150bp) that overlapped with known Pho binding sites identified by Kassis and colleagues (Fig. 4C). Importantly, these regions correspond well with PREs predicted by modENCODE (Fig. 4C). As shown in Fig. 4D, this analysis identified 436 putative PREs in the *H3.3*^*WT*^*H3.2*^*HWT*^ and 189 regions in the *H3.3A*^*null*^*H3.2*^*HWT*^ animals. We further narrowed this list to the common set of 125 putative PREs that were identified independently in both control genotypes (Fig. 4D).

To ascertain whether the K36R mutation impedes spreading of H3K27me3, we generated heatmaps and metaplots of H3K27me3 signal in the regions flanking these presumptive PREs (Fig. 4E). Although the short-fragment reads were useful for identifying regions likely to harbor DNA binding proteins at PREs, we used reads from pooled bigwig files of supranucleosomal fragment length (150-700bp) to compare H3K27me3 signal in the regions +20kb flanking the presumptive PREs for each genotype. Strikingly, metaplots for the two control genotypes, as well as the *H3.3*^*K36R*^*H3.2*^*HWT*^ mutant were nearly identical, demonstrating that mutation of H3.3K36 alone had no effect on spreading across broad Pc domains (Fig. 4E). In contrast, the *H3.3*^*WT*^*H3.2*^*K36R*^ mutant displayed a marked depletion of H3K27me3 signal in the flanking regions, but not directly over the PREs themselves (Fig. 4E). These data not only confirm and extend previous findings (*27*) but also demonstrate that *in vivo*, H3.2K36 plays a much more important role than H3.3K36 in facilitating spreading of H3K27me3 across broad domains, suggesting that H3.3K36 promotes proper Hox gene regulation by a different means.

### Loss of H3.3K36 has no effect on H3K27 trimethylation at PREs

Previous reports in mouse embryonic stem cells and *Drosophila* showed that H3.3 is enriched at CpG islands and PREs, regions that most consistently function as PRC2 nucleation sites, respectively (*61, 62*). Furthermore, GAGA factor (GAF), which can bind PREs and promote repression of associated genes (*63-66*), interacts with HIRA, an H3.3-specific histone chaperone (*67*). Although the analysis of long-fragment reads in Fig. 4 failed to identify a loss of signal within broad domains, we wondered if the *H3.3*^*K36R*^*H3.2*^*HWT*^ animals might display a more localized effect at PREs.

In CUT&RUN, the distribution and overall signal from different fragment sizes are functions of both the relative presence of the epitope of interest and the relative accessibility of the surrounding chromatin (Fig. 5A, (*52*)). To interrogate the H3K27me3 profiling data more specifically, we generated meta-plots +500bp around 344 short fragment peak summits from all four genotypes that overlapped with the set of robust PREs established in Figure 4. As shown in Fig. 5B, we observed no appreciable difference in signal between the *H3.3*^*K36R*^*H3.2*^*HWT*^ mutant and its control (Fig. 5B). Despite reported colocalization of H3.2 and H3.3 at PREs in S2 cells (*62*), we also saw no effect at PREs in the *H3.3*^*WT*^*H3.2*^*K36R*^ mutant (Fig. S10). These data show that individual loss of H3.2K36 or H3.3K36 has little effect on H3K27 methylation immediately flanking presumptive PREs.

Because PREs are heterogeneous with respect to DNA sequence, binding factor occupancy, and functional properties (*32, 59, 68*), we wondered if certain PREs might be preferentially affected by the H3.3K36 mutation even though we detected no general effect. To address this question, we performed DESeq2 differential expression analysis on short fragment read counts from the K27me3 pellet fraction in all four genotypes (Fig. 5C). We broadened the approach to examine intervals that overlapped with a more general set of short fragment peak summits as well as with Pho binding sites (Figs. 5C, S10). Remarkably, we identified no differential peaks (0/711) overlapping with Pho sites in the *H3.3*^*K36R*^*H3.2*^*HWT*^ mutant (Fig. 5C). Similarly, there were only a few differential peaks (41/711) in the *H3.3*^*WT*^*H3.2*^*K36R*^ mutant (Fig. S10). These data show that mutation of H3.2 or H3.3 lysine 36 has little effect on the accessibility of PREs to MNase cutting or on H3K27 trimethylation of nucleosomes that flank Pho binding regions.

### H3.3 is enriched at Pho binding sites inside and outside of Pc domains

Chromatin profiling for H3K27me3 in the *H3.3*^*WT*^*H3.2*^*K36R*^ mutant supports a clear role for H3.2 in propagation of H3K27me3 between PREs (Fig. 4). Given that H3.3 is reportedly enriched at PREs in S2 cells (*62*), the failure to observe a reduction in H3K27me3 signal at these sites in *H3.3*^*K36R*^*H3.2*^*HWT*^ mutant wing discs was surprising (Fig. 5B-C). We therefore determined whether H3.3 is also enriched at PREs in the wing disc by CUT&RUN profiling using anti-H3.3 antibodies on *yw* (positive) and *H3.3Δ* mutant (negative) control genotypes. Libraries were sequenced and normalized as described above and in the Methods. As shown in Figure 5D-F, analysis of the anti-H3.3 and -H3K27me3 CUT&RUN data alongside sets of previously validated PREs (*54*) and Pho binding sites (*59*) revealed a number of interesting features.

We detected an enrichment of H3.3 at Pho-positive PREs located in the Bithorax complex as well as in other H3K27me3 domains (Fig. 5D). Note that the presence of H3.3 at these was obvious when visualized on a genome browser (high signal, low noise) but the peak summit intensities often did not meet the threshold of the MACS2 peak caller. Consistent with the notion that H3.3 is enriched within active chromatin (*61, 69-71*), we observed much higher levels of H3.3 in “euchromatic” regions located outside of repressive H3K27me3 domains (Fig. 5D). Interestingly, the H3.3 peaks in euchromatic regions also coincided with Pho binding sites, and visual inspection suggested they were typically of greater intensity than those located inside of H3K27me3 domains (Fig. 5D).

To determine the significance of the association between H3.3 and Pho sites, we performed Fisher’s Exact Test on intervals known to have Pho binding potential (*59*) and those enriched for H3.3. As shown in Fig. 5E, the correlation between H3.3 and Pho peaks was highly significant, regardless of their H3K27me3 status (p < 7.06 × 10^−108^ inside; p < 2.52 × 10^−213^ outside). To quantify the difference in H3.3 signal at Pho sites inside vs. those outside of Pc domains, we generated heatmaps and metaplots for H3.3 signal at sites of overlapping H3.3 and Pho binding potential (Fig. 5F). We found that H3.3 enrichment was ∼40% greater at Pho sites outside K27me3 domains compared to those inside (Fig. 5F). We note that the H3.3 peaks within K27me3 domains were often too shallow to be identified by MACS2; thus the relative degree of enrichment outside of K27me3 domains is likely an underestimate. Taken together, our data show that H3.3 is indeed enriched at wing disc PREs; however, mutation of H3.2 or H3.3 K36 has very little effect on H3K27me3 signal at these sites.

### Complete loss of H3K36 results in profound developmental defects

Our genetic experiments demonstrate that both H3.2K36 and H3.3K36 contribute to accurate repression of Hox genes. However, Western blotting and CUT&RUN experiments suggest that they contribute to this process by different mechanisms. While *H3.3*^*WT*^*H3.2*^*K36R*^ animals clearly have a defect in H3K27 trimethylation, *H3.3*^*K36R*^*H3.2*^*HWT*^ mutants do not. Furthermore, both H3.2K36R and H3.3K36R mutant animals exhibit relatively mild homeotic phenotypes compared to animals with mutations in other genes required for this process (*43, 44, 72, 73*).

To investigate potential redundancy between H3.3K36 and H3.2K36, we engineered a combined *H3.3*^*K36R*^*H3.2*^*K36R*^ mutant (Fig. S4). When H3 lysine 36 is completely removed from the genome (*H3.3*^*K36R*^*H3.2*^*K36R*^), we observed a much stronger developmental phenotype than either the H3.2K36R or H3.3K36R alone, as none of the *H3.3*^*K36R*^*H3.2*^*K36R*^ progressed beyond the first larval instar (L1) stage. *H3.3*^*K36R*^*H3.2*^*K36R*^ mutant embryos frequently exhibited gross morphological defects, such as the ventral nerve cord and gut defects (Fig. 6A-B). Indeed, only ∼30 % of the *H3.3*^*K36R*^*H3.2*^*K36R*^ embryos hatched (Fig. 6C). Thus, while H3.3K36 is not required to complete development, it compensates to a large degree for loss of H3.2K36. Taken together, the results reveal that H3K36 is critical for proper embryogenesis, and that H3.2K36 and H3.3K36 can compensate for one another during these early stages of development.

**Figure 6.**
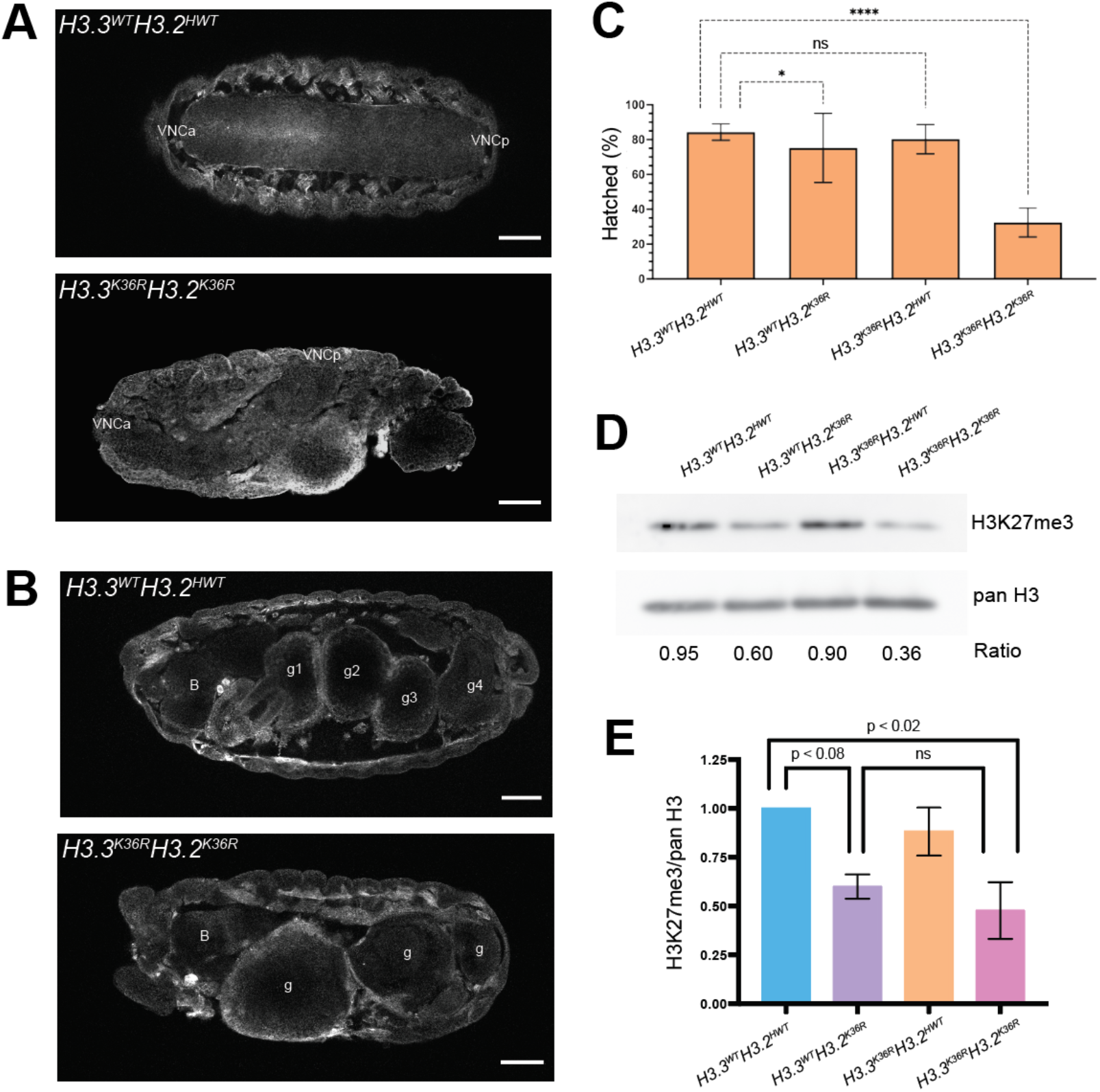
Analysis of combined *H3.3*^*K36R*^*H3.2*^*K36R*^ mutants during early development. **(A)** Stage 16 embryos of *H3.3*^*K36R*^*H3.2*^*K36R*^ mutants and *H3.3*^*WT*^*H3.2*^*HWT*^ controls were fixed and stained with an anti-GFP antibody, detecting YFP protein, to outline gross morphology. The top panel displays a control animal with a straight and symmetrical ventral nerve cord (VNC). Anterior and posterior VNC is designated (VNCa, VNCp). The bottom panel displays a mutant embryo exhibiting severe VNC defects. Overall, we observed a similar degree of VNC twisting in 4/44 (∼9%) *H3.3*^*K36R*^*H3.2*^*K36R*^ mutant embryos, and 0/73 instances in controls. This phenotype was never observed in other H3K36R mutant genotypes where we examined embryos. **(B)** Embryos, as in A, but highlighting severe defects in gut development. B = brain; g1, g2, g3, g4 correspond to the 4 gut lobes characteristic of this stage. We observed several examples of this phenotype in mutants, but zero in controls or other mutant genotypes. For panels A and B, scale bar = 50µm. Embryos are displayed with anterior at the left, posterior to the right. **(C)** Embryonic viability assay (n=250-400 embryos). The fraction of embryos progressing from embryonic to L1 stages (%Hatched) was calculated. For the *H3.3*^*WT*^*H3.2*^*K36R*^ and *H3.3*^*K36R*^*H3.2*^*K36R*^ genotypes, this value also reflects an adjustment for the presence of a balancer chromosome in 50% of the embryos. Statistical significance was calculated with GraphPad Prism software using Fisher’s exact test independently comparing each H3K36R mutant genotype with the *H3.3*^*WT*^*H3.2*^*HWT*^ control. P-values are abbreviated as in Figure 1. **(D)** Western blotting using anti-H3K27me3 and -panH3 antibodies was performed from extracts of first instar (L1) mutant and control larvae. **(E)** Quantification of four independent replicates is shown. Mean pixel intensity in equal sized boxes was calculated for each band, and a ratio of H3K27me3/panH3 was calculated for each biological replicate. For each experiment, the H3K27me3/panH3 ratio of mutant genotypes was normalized to *H3.3*^*WT*^*H3.2*^*HWT*^ controls. For each genotype, we plotted the normalized mean and SD of H3K27me3/panH3 values of all four replicates. Statistical significance between raw H3K27me3/H3 ratios was assessed by paired one way ANOVA, followed by Friedman Tests individually comparing mutant genotypes to *H3.3*^*WT*^*H3.2*^*HWT*^ controls. P-values are directly noted on the graph. *H3.3*^*K36R*^*H3.2*^*HWT*^ mutants are no different from *H3.3*^*WT*^*H3.2*^*HWT*^ controls (p-value = ns, not shown on graph), whereas, *H3.3*^*WT*^*H3.2*^*K36R*^ and *H3.3*^*K36R*^*H3.2*^*K36R*^ exhibit a sharp decrease in H3K27me3 levels.

### Total loss of H3K36 does not alter global H3K27me3 levels but causes synergistic misexpression of Hox proteins

Because H3K27 trimethylation at PREs is believed to be a prerequisite for the spread of this PTM into flanking regions (reviewed in (*74*)), one might expect that if H3.2 and H3.3 functioned redundantly to enable PRC2 activity at PREs, levels of H3K27me3 in the combined *H3.3*^*K36R*^*H3.2*^*K36R*^ would be further diminished compared to what is observed in *H3.3*^*WT*^*H3.2*^*K36R*^ animals. To test this idea, we performed western blotting of first instar (L1) lysates from *H3.3*^*WT*^*H3.2*^*HWT*^ (control), *H3.3*^*WT*^*H3.2*^*K36R*^, *H3.3*^*K36R*^*H3.2*^*HWT*^, and *H3.3*^*K36R*^*H3.2*^*K36R*^ genotypes using anti-H3K27me3 antibodies, along with a pan-H3 loading control (Fig. 6D-E). In total, we performed and quantified four biological replicates. In the *H3.3*^*WT*^*H3.2*^*K36R*^ genotype, we observed a ∼40% reduction (p < 0.08) in H3K27me3 compared to the *H3.3*^*WT*^*H3.2*^*HWT*^ control (Fig. 6D-E). This finding is consistent with a previous report showing similarly reduced H3K27me3 levels in wandering third instar *H3.3*^*WT*^*H3.2*^*K36R*^ larvae (*12, 27*). As expected, we observed no change in H3K27me3 levels in the *H3.3*^*K36R*^*H3.2*^*HWT*^ mutants. Notably, we observed no significant change H3K27me3 in the combined *H3.3*^*K36R*^*H3.2*^*K36R*^ genotype compared to the *H3.3*^*WT*^*H3.2*^*K36R*^ mutant, with H3K27me3 at ∼40% of the *H3.3*^*WT*^*H3.2*^*HWT*^ control (Figs. 6D-E). These data suggest that H3.2 and H3.3 K36R mutations are unlikely to function redundantly at PREs by directly inhibiting PRC2.

Mutation of either residue alone confers relatively mild defects in Hox gene expression and homeotic transformation. To further explore the possibility of genetic redundancy between H3.2K36 and H3.3K36, we carried out immunostaining of control, single and combined K36R mutations and determined the extent to which Hox proteins were misexpressed. In wild type embryos, AbdB is limited to embryonic parasegments PS10-PS14, with the highest expression levels near the posterior of this region. These parasegments roughly correspond to adult abdominal segments A4-A9 (*75*). Immunostaining of Stage 16 *H3.3*^*WT*^*H3.2*^*HWT*^ embryos recapitulated the established pattern of AbdB expression in the ventral nerve cord (VNC, Fig. 7A, bracket). We also observed the mild, stochastic derepression of AbdB in *H3.3*^*WT*^*H3.2*^*K36R*^ embryos noted previously (*27*), although the penetrance of this phenotype was incomplete (Fig. 7A, arrowheads). The *H3.3*^*K36R*^*H3.2*^*HWT*^ phenotype was similar to that of the control, in that we never observed AbdB derepression in anterior parasegments (Fig. 7A). In contrast, *H3.3*^*K36R*^*H3.2*^*K36R*^ mutants displayed extensive derepression of AbdB in anterior parasegments, comparable to that of *H3.3*^*WT*^*H3.2*^*K27R*^ mutants (Fig. 7A, arrowheads). We conclude that H3.3K36 and H3.2K36 can functionally compensate for one another to fully repress AbdB expression in the embryonic VNC.

**Figure 7.**
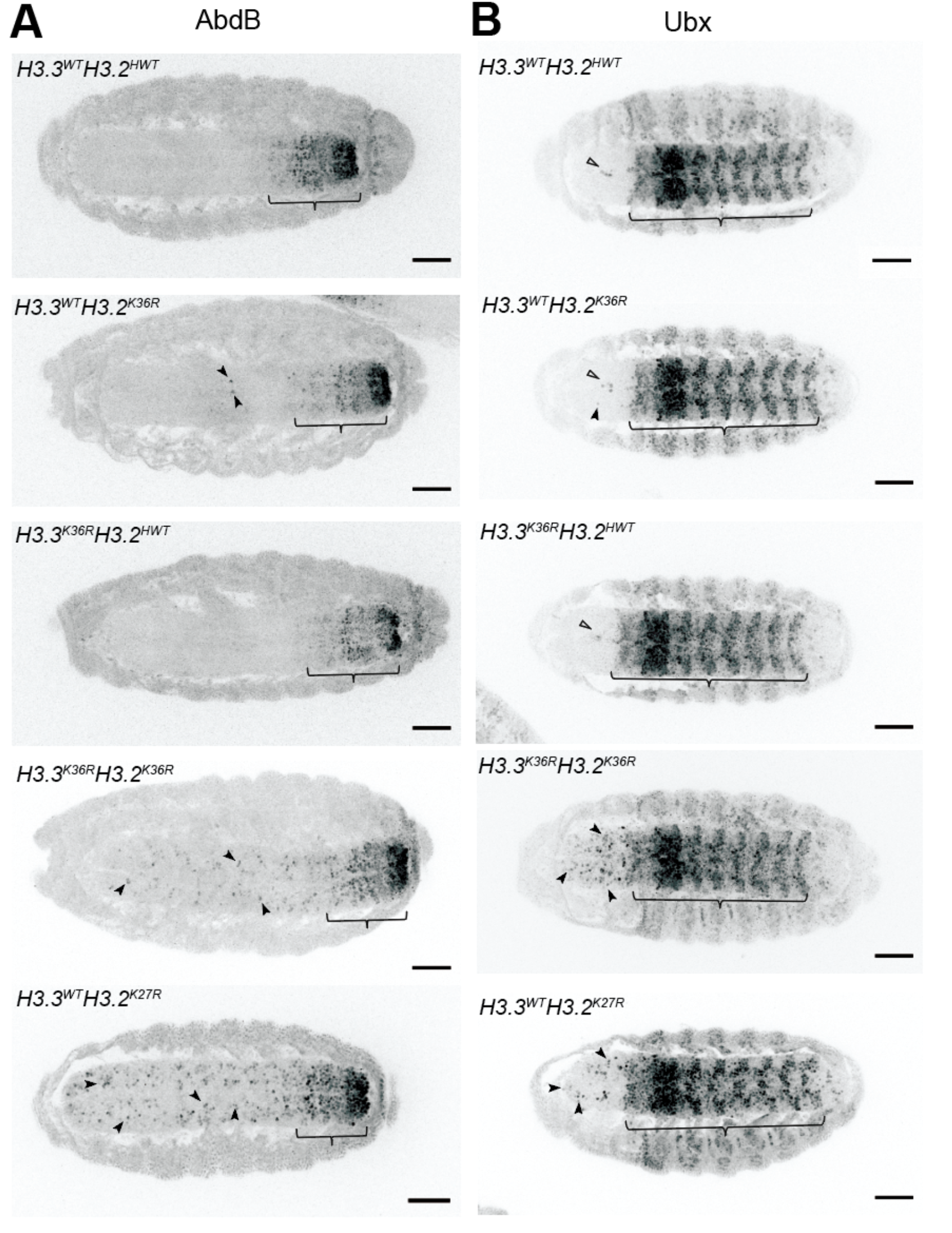
Combined mutation of *H3.3* and *H3.2 lysine 36* causes synergistic derepression of Hox genes. **(A)** Stage 16 embryos of various histone H3 mutant genotypes and controls were fixed and stained with anti-AbdB antibodies. Embryos were stained with anti-GFP antibodies to detect YFP for staging and genotype selection. Images used for staging these embryos can be found in Fig. S12. Brackets indicate the expected boundary of AbdB expression in wild type embryos. Filled arrows highlight individual cells exhibiting anterior derepression of AbdB. Scale bar = 50µm. Overall, the individual *H3.3*^*WT*^*H3.2*^*K36R*^ and *H3.3*^*K36R*^*H3.2*^*HWT*^ mutants closely resemble the *H3.3*^*WT*^*H3.2*^*HWT*^ negative controls, whereas the double mutant *H3.3*^*K36R*^*H3.2*^*K36R*^ embryos are like the *H3.3*^*WT*^*H3.2*^*K27R*^ positive control. **(B)** Same as panel A, except stained with anti-Ubx. Empty arrows highlight a small patch of Ubx positive cells consistently observed in wild type controls, and thus do not indicate H3K36 dependent derepression.

To further assess redundancies between H3.2K36 and H3.3K36 in Hox gene repression, we analyzed expression of another Hox protein, Ubx. In Stage 16 embryos, control *H3.3*^*WT*^*H3.2*^*HWT*^ embryos recapitulated the known Ubx staining pattern (*75*), though we consistently observed a small cluster of Ubx expressing cells at the midline, slightly anterior to the known boundary at PS5 (Fig. 7B, arrowheads). Because these Ubx-positive cells were always present in the controls, we did not score them as derepression events in any mutant genotype. Notably, whereas Finogenova et al. (*27*) observed ectopic expression of Ubx in *H3.3*^*WT*^*H3.2*^*K36R*^ third instar (L3) wing discs, they failed to detect derepression of Ubx in Stage 16 embryos (*27*). Likewise, *H3.3*^*WT*^*H3.2*^*K36R*^ and *H3.3*^*K36R*^*H3.2*^*HWT*^ embryos resembled the *H3.3*^*WT*^*H3.2*^*HWT*^ control. In contrast, we found extensive Ubx expression in anterior parasegments of the VNC of *H3.3*^*K36R*^*H3.2*^*K36R*^ embryos which was comparable to that of *H3.3*^*WT*^*H3.2*^*K27R*^ embryos. As with AbdB, we conclude that H3.2K36 and H3.3K36 can functionally compensate for each other in the repression of Ubx expression in the embryonic VNC.

Together with *abdA*, the *AbdB* and *Ubx* genes are clustered within the Bithorax Complex (BX-C). In addition to BX-C, the Antennapedia Complex (ANT-C) is located roughly 10Mb away, and contains the *Antp, Scr, Dfd, pb*, and *lab* genes (*76, 77*). We also performed immunostaining for the Antp protein (Fig. S13). As with AbdB and Ubx, we also observed incompletely penetrant stochastic derepression of Antp in *H3.3*^*WT*^*H3.2*^*K36R*^ embryos, wild type staining in the *H3.3*^*K36R*^*H3.2*^*HWT*^ embryos, and more extensive and penetrant derepression in *H3.3*^*K36R*^*H3.2*^*K36R*^ mutant embryos. In general, Antp derepression in *H3.3*^*K36R*^*H3.2*^*K36R*^ was less extensive than that of *H3.3*^*WT*^*H3.2*^*K27R*^ embryos. These results reveal that H3.2K36 and H3.3K36 can also functionally compensate for one another to carry out repression of *Antp*. Taken together, the data are consistent with the hypothesis that lysine 36 of H3 is necessary for maintaining Hox gene repression *in vivo*, and that the ability of H3K36 to enable this repression can be provided by either H3.2 or H3.3 (*12, 27*).

## Discussion

This study demonstrates that H3.2K36 and H3.3K36 collectively mediate proper repression of Hox genes throughout development. At embryonic stages, combined mutation of these two H3 variants causes profound dysregulation of Hox genes. Later in development, the individual *H3.2* and *H3.3* K36R mutants each exhibit mild homeotic transformations and enhance PcG homeotic phenotypes. Yet our study strongly suggests that H3.2K36 and H3.3K36 promote proper Hox gene repression by distinct mechanisms.

Previous studies suggested a critical role for unmodified H3.2K36 in E(z) mediated H3K27 trimethylation (*27*). This work corroborates and extends those observations, demonstrating that H3.2K36 is critical for maintaining H3K27me3 across numerous Pc domains genome-wide. We also show *in vivo* that this role in sustaining PRC2 function is specific to H3.2, as we observed no change in global levels or distribution of H3K27me3 in *H3.3*^*K36R*^ mutants, and there was no additional impact when combined with an *H3.2*^*K36R*^ mutation. Even in a sensitized Pc^3^/+ genetic background, where *H3.3*^*K36R*^ mutants exhibit extensive homeotic transformations, global H3K27me3 levels are not significantly changed. In addition to having no effect on the overall distribution of H3K27me3, *H3.3*^*K36R*^ mutation also elicits no specific effect at PREs (Fig. 5B-C). Perhaps surprisingly, the same is true for *H3.2*^*K36R*^ mutation. Despite a clear defect in the *H3.2*^*K36R*^ mutant in inter-PRE regions, H3K27me3 levels are the same as the control at putative PREs. Moreover, western blotting suggests that global H3K27me3 levels are not significantly reduced when both *H3.2* and *H3.3* K36R mutations are combined. To more definitively address the possibility that localized loss of H3K27 trimethylation is responsible for Hox gene derepression, chromatin profiling of the combined mutant in L1 larvae from a relatively homogeneous cell population would be necessary. Such an experiment presents numerous technical hurdles, but may be possible in the future.

Another possibility for why loss of H3K36 might fail to affect H3K27me3 levels at PREs could be because PRC2 activity is greatly enhanced at these locations by other factors. Indeed, vertebrate Polycomb-like (PCL) proteins interact with H3K36me3 (*78-80*), and function as enhancers of PRC2, and EZH2 activity (*81-83*). Notably, structural analysis of the fly ortholog (Polycomblike, Pcl), suggests that this protein is unlikely to bind H3K36me3 (*84*). However, Pcl retains the ability to bind E(z) in the context of a large complex containing PRC2 (*85, 86*), and *Pcl* mutants exhibit reduced H3K27me3 (*87*). Future studies addressing the role of Pcl at *Drosophila* PREs in the context of H3K36R mutations will be of great interest.

During early stages of development, we have shown that arginine substitutions in histone H3.3 at either K27 (*39*) or at K36 (this work) have little effect on overall levels of H3K27 trimethylation. Similarly, overexpression of H3.3K36R histones in cultured human cells had no effect on H3K27me3 (*47*). These results are consistent with a model put forth by the Allis laboratory, suggesting that H3.3 does not provide a methyl-substrate nucleosome at PREs. Instead, increased turnover of H3.3-containing nucleosomes is thought to be a key feature in PRC2 recruitment to PREs (*88*). By extension, higher nucleosome turnover at PREs would be more important for *establishing* a silent chromatin domain than it would be for *maintaining* one. Thus, our finding that H3.3 occupancy was much higher at Pho sites located outside of H3K27me3 domains compared to those inside (Fig. 5E-F) supports this model.

Given that PREs are thought to be multifaceted regulatory modules that can function as repressors or enhancers in different developmental contexts (*54, 89, 90*), it is intriguing to speculate on other roles that H3.3K36 might play in Hox gene regulation. Our earlier work demonstrates that *H3.2*^*K36R*^ mutants exhibit widespread transcriptomic defects (*12*), and mutants for all three H3K36 lysine methyltransferases in *Drosophila* also produce large changes in gene expression in the larval brain (*91*). Given the enrichment of H3.3 in active areas of the genome, one would expect similar dysregulation in *H3.3*^*K36R*^ mutants. Therefore, it is possible that H3.3K36 is key to maintaining proper expression levels of one or more genes involved in Pc gene silencing. Transcriptomic studies in the *H3.3*^*K36R*^, *H3.2*^*K36R*^ and combined mutants would address this possibility.

A role in 3D genome organization is another potential mechanism for how H3.3K36 might promote Pc silencing. Several lines of evidence point to the connection between Pc domains, H3K36me, and genome organization (*92, 93*). Furthermore, foci of PcG proteins bound at different chromosomal sites coalesce in “Polycomb Bodies” (*94-96*). Contacts between Hox gene clusters appears to be functionally important to gene silencing, as mutation of the Fab-7 element in the BX-C, which strengthens association between the two HOX gene clusters, weakens silencing of genes in the ANTP complex (*94*). Notably, 3D contacts within Pc domains correspond preferentially to PREs, where H3.3 is enriched (*62*). In mouse, knockdown of H3.3 (but not H3.1), leads to increased chromatin compaction (*97, 98*). Thus, studies probing the effect of an H3.3K36 mutation on 3D genome organization could also be of great interest.

Despite several intriguing findings and conclusions, our histone gene replacement approach has several limitations. The K36R mutation effectively abolishes all K36 PTMs, including acetylation and individual states of lysine methylation (me0, me1, me2, and me3). The simultaneous loss of me0 and me2 is particularly relevant to a discussion of Pc target loci. The Ash1 methyltransferase dimethylates H3K36 (*26, 99, 100*). In *Drosophila*, mutation of Ash1 impairs the expression of Hox genes, including *abdB* and *Ubx*, in compartments where they are normally expressed (*28, 101*), and *in vitro* studies confirm that the H3K36me2 mark inhibits E(z) function (*26, 101*). In this context, an H3K36R mutation is analogous to an *ash1/E(z*) double mutant. Previous studies showed that *ash1*/*E(z*) double mutant clones exhibited an Hox gene derepression phenotype similar to that of an *E(z*) single mutation (*102*). Thus, *E(z*) is epistatic to *ash1*. This observation is consistent with our results in the combined K36R mutant embryos.

In summary, this study implicates the lysine 36 residue of non-centromeric H3 variants in promoting Hox gene repression in flies. Yet despite their nearly identical amino acid sequence and potential to permit E(z) mediated H3K27me3, they contribute to Hox gene repression by non-identical means. Future work should elucidate the roles played by H3K36 PTMs in different parts of the genome, how they contribute to chromatin organization, and in the various steps of pre-mRNA processing.

## Materials and Methods

### Drosophila Lines and Husbandry

To obtain progeny of crosses, parental flies were housed in cages plugged with grape juice agar plates containing supplemental yeast paste. Plates were changed at least daily. Embryos and L1 larvae were harvested directly from grape juice plates. Older animals were handpicked at 2^nd^ instar stage, 50 per vial, and raised on standard cornmeal-molasses food. All experimental animals were raised at 25°C.

Details regarding construction of bacterial artificial chromosome (BAC) transgenes containing the 12xH3.2 histone gene arrays can be found here (*10*). The 12x *HWT, K27R, K36R, K9R* alleles were generated previously (*9, 13*). Flies carrying the *HisΔ, twGal4* and *HisΔ, UAS:2xYFP* chromosomes (*103*) were gifts from A. Herzig. The *H3.3A*^*2×1*^ (*H3.3A*^*null*^) and *H3.3B*^*0*^ (*31*), along with the *Df(2L)Bsc110* deficiency and *Pc*^*3*^ allele were obtained from Bloomington Stock Center (#68240, #8835, and #1730).

### Generation of Mutant Genotypes

For detailed crossing schemes, see Fig. S1. Animals of the *HisΔ* genotype were obtained by selection for YFP. Other H3.3 genotypes were selected for absence of a *CyO, twGFP* balancer chromosome. When not universal in progeny, selection for *12xH3.2* transgenes were identified in embryos by selecting animals that progressed beyond cellularization, and by picking YFP positive larvae at later stages. Adults scored for homeotic transformations were selected by the presence or absence of adult body markers.

### CRISPR/Cas9 Mutagenesis of H3.3B Locus

Cas9-mediated homologous recombination of H3.3B was carried out as described previously (*14, 39*), with the following differences. A single guide RNA targeting the *H3.3B* gene near the K36 residue was inserted into pCFD3 and co-injected along with a 2-kb repair template containing the H3.3BK36R substitution. Constructs were injected into embryos expressing Cas9 from the *nanos* promoter (*nanos-cas9*; (*104*)). Recovered *H3.3B*^*K36R*^ alleles were subsequently crossed into a *H3.3A* null background [*H3.3A*^*2×1*^ over deficiency *Df*(*2L*)*BSC110*].

### Pupal and Adult Viability Assays

For each genotype, fifty 2^nd^ instar larvae were picked from grape juice agar plates and transferred to vials containing molasses-cornmeal food. Pupal cases and eclosed adults were counted until 13 and 18 days post egg-laying, respectively. Pupal and adult viability percentages were calculated on a per vial basis by dividing the number of pupal cases or eclosed adults per 50 input larvae, and multiplying by 100. Each vial constituted one biological replicate for statistical purposes. Between 200-400 total animals were analyzed per genotype.

### Embryonic Hatching Assay

For each genotype, *HisΔ*, YFP+ embryos were moved onto a clean grape juice plate with a pick into lines of 50. At Day1 and Day2 post selection, empty eggshells were counted and recorded as “hatched”. Hatching frequency (%) was calculated by dividing the total number of empty eggshells after 2 days by either the number of total embryos (for *12xH3.2*^*HWT*^ genotypes), or by 0.5x the total number of embryos (for *12xH3.2*^*K36R*^ genotypes), and multiplying by 100. The 0.5x adjustment is due to the necessity of carrying the *12xH3.2*^*K36R*^ transgene over *TM6B*, and the inability to distinguish those two genotypes at the embryonic stage. Between 250-400 animals were analyzed per genotype.

### Immunofluorescence

Embryos were collected for 3 hours on grape juice agar plates and then aged for 12 hours at 25°C. Embryos of appropriate age were collected in mesh baskets, dechorionated for 5 minutes in 50% bleach, and rinsed thoroughly in deionized water, followed by Embryo Wash Buffer (EWB: 1X PBS, 0,03% Triton-X). Embryos were transferred to a glass vial, where first 0.5 ml of fixative (1X PBS, 4% formaldehyde) was added, followed by 0.5ml heptane, and nutated for 20 minutes. The bottom layer of fixative was completely removed and replaced with 0.5ml 100% methanol. Embryos were devitellinized by vigorous shaking for 30 seconds. After removal of methanol and heptane, embryos were washed for 5 minutes, twice with 100% methanol, once with 1:1 methanol:PBS-T (1X PBS, 0.15% Triton-X 100), twice with 1X PBS-T. This was followed by two additional 30 minute PBS-T washes. All washes were performed with nutation during the incubation period. Between the two 100% methanol washes, samples were transferred from the glass vial to an Eppendorf tube.

Embryos were blocked in PBT with 2% normal goat serum (NGS) for one hour at RT. Primary antibody incubations were performed overnight at 4°C in PBS-T with 2% NGS plus one or more of the following antibodies: polyclonal rabbit anti-GFP (1:800, Abcam #ab290), monoclonal mouse anti-Abd-B (1:500, Developmental Studies Hybridoma Bank #1A2E9), monoclonal mouse anti-Ubx (1:500, Developmental Studies Hybridoma Bank #FP3.38), monoclonal mouse anti-ANTP (1:250, Developmental Studies Hybridoma Bank #4C3). Embryos were washed 3x for 10min in PBS-T. Secondary antibody incubations were performed for 45 minutes at RT with both anti-rabbit-Alexa Fluor-546 (Invitrogen #A11035) and anti-mouse Alexa Fluor-488 (Invitrogen #A21202) in PBS-T, followed by 2min DAPI stain, and 3 additional PBS-T washes. Embryos were mounted in Vectashield® mounting media and imaged on a Leica SP8 Confocal Microscope, 20x oil immersion objective. Images were viewed and analyzed using ImageJ.

### Western Blotting

Protein lysates from 1^st^ instar larvae or adult heads were obtained by homogenization with a micropestle in SUTEB buffer (1% SDS, 8M Urea, 10mM EDTA, ph 8.0, 5% beta-mercaptoethanol (BME), with 1:20 Halt Protease Inhibitor Cocktail (Thermofisher #78429)). Chromatin was further disrupted by sonication with the Bioruptor Pico (Diagenode).

Samples were run on 4-15% Mini-PROTEAN® TGX Stain-Free™ Protein Gels (Bio-Rad #4568084) for 60 minutes at 100V. Western blotting was performed using the Biorad Trans Blot Turbo transfer system using the provided buffer (Biorad #10026938) onto a nitrocellulose membrane at 1.3A/25V for 7 minutes. Membranes were blocked at RT in 5% milk in TBS-T. Primary antibody incubation occurred overnight at 4°C in TBS-T with 5% milk with one of the following antibodies: polyclonal rabbit anti-H3K27me3 (Activemotif #39055, 1:1000), polyclonal rabbit anti-H3 (Abcam #1791). For each primary antibody, an anti-Rabbit secondary (Sigma #12-348) was used at 1:5000.) Blots were incubated with chemiluminescent detection reagent (Amersham ECL™ Prime Western Blotting Detection Reagents, GE Healthcare #RPN2236) and imaged on an Amersham Imager 600 (GE Healthcare). Between primary antibody antibodies, blots were stripped, rinsed in TBS-T, and incubated with detection reagent to verify removal of antibody before reprobing.

Relative band intensity ratios were calculated on ImageJ. Briefly, a box of equal size and dimension was drawn around each band and integrated density (IntDen) inside the box was recorded. For each blot, a ratio of H3K27me3/H3 IntDen was calculated per sample lane. Additionally, a normalized value for each mutant genotype was calculated by dividing the mutant ratio by that of the *H3.3*^*WT*^*H3.2*^*HWT*^ genotype (for L1 experiments) or by individual control indicated in Figure legend (adult heads).

### CUT&RUN Chromatin Profiling

We performed CUT&RUN in wing disc tissue as described in (*105*), modified from (*49*). For each replicate, twenty WL3 imaginal wing discs along with two wing discs of *D.yakuba* were used per replicate. For the H3K27me3 experiment, αH3K27(CST#9733,1:100) and protein AG-MNase (1:100, UNC core) were used. For the H3.3 experiment, αH3.3 (H3F33B, Abnova #H00003021-M01, 1:100) and protein A-MNase from the Henikoff lab were used

The Thruplex DNA-seq kit was used for the library preparation. The manufacturer’s protocol was followed until the amplification step. After the addition of indexes, 16-21 cycles of 98C, 20s; 67C, 10s were run. DNA library purification was done using AMPure XP beads. Libraries were sequenced on an Illumina NextSeqP2.

### Bioinformatic Analyses

A flowchart with details of our bioinformatic workflow can be found in Fig.S6. CUT&RUN data were processed as in https://github.com/mckaylabunc/cutNrun-pipeline with minor modifications. Differential peak analysis was performed using featureCounts (*106*) and DESeq2, v1.34.0 (*107*). Details for how each list of peaks or intervals for analysis was determined can be found in Fig. S8. For broad domains (Fig. 4B), pellet reads of all fragment sizes were used for DESeq2 analysis. For PRE based analyses in Figs. 5C, S10B, and S11, only short fragment (20-120bp) pellet reads were analyzed with DESeq2.

All heatmaps and metaplots were generated with deepTools, v3.5.1 (*108*). The Y chromosome was excluded from analysis. For details of how reference points were obtained for each analysis, see Fig. S8. For broad domain analysis (Fig. 4D), deepTools, v3.5.1 was used to calculate signal intensity from K27me3 z-score normalized pellet large fragment pooled bigWig files at “robust putative PREs” (See Fig. S8), and for 20kb flanking these regions for heat maps and metaplots. For PRE based heatmaps and metaplots in Figs. 5B and S10A, deepTools, v.3.5.1 was used to calculate and plot signal intensity from K27me3 z-score normalized pellet large fragment and small fragment pooled bigWig files at these “concatenated PRE intervals” and for 500bp flanking regions. For Fig. 5F heatmaps and metaplots, deepTools, v.3.5.1 was used to calculate signal intensity from H3.3 z-score normalized supernatant all fragment pooled *yw* bigWig files at Pho binding sites (see Fig. S8) and 5kb flanking regions inside and outside broad H3K27me3 domains.

### Statistical Analysis

Statistical analyses for non-genomic experiments were performed and guided by GraphPad Prism according to the characteristics of each data set. Details of each individual statistical analysis can be found on each corresponding figure legend. Adjusted p-values obtained to call differential peaks in the CUT&RUN experiment were determined with the DESeq2 analysis package (*107*).

The 2-tailed Fisher’s Exact Test in Fig. 5E, was performed between Pho binding intervals and H3.3 MACS2 narrow peak summits + 150bp intervals from the *yw* genotype with *H3.3Δ* peaks blacklisted, separately by H3K27me3 status using BEDtools, v.2.3.0 (*109*).

### Homeotic Transformation Assay

Individual crosses for each genotype (see Fig. S3 for details) were set up in cages, fed daily with grape juice agar and yeast paste. Groups of fifty 2^nd^ instar larvae were collected per vial of cornmeal molasses food where they grew to adulthood. Adult males of the correct genotype were separated by body markers and scored for homeotic transformations under a Leica M60 stereomicroscope. Images were obtained using an iPhone camera mount on the Leica M60.

### Scanning Electron Microscopy

Flies collected and stored in 70% ethanol were sequentially deyhydrated in 100% ethanol. Fly bodies were mounted, and images collected using a Hitachi 392 TM4000Plus tabletop SEM microscope using 250x magnification.

### Data and materials availability

Raw and processed RNA-seq data were deposited to Gene Expression Omnibus (GEO) under accession GSE215017. The accession is currently private until manuscript acceptance. BED files from the following previously published ChIP Seq data sets (GSE102339) were used (*59*). BED files derived from Pho_WT_1 (GSM2634951), Pho_WT_2 (GSM2634952), and Pho_WT_3 (GSM2634953) were also used. All other data needed to assess the conclusions in this paper are included in the main paper and/or Supplementary Materials. Additional information available from the corresponding author on request.

## Acknowledgments

We thank the Bioinformatics Analytics Research Collaborative (BARC) at the University of North Carolina (UNC) School of Medicine for facilitating access to bioinformatics education and training (for HRS and BDM) as well as staff support (for JMS). Additional educational support was provided by the UNC Chancellor’s Science Scholars program (to KMA) and by the UNC SOLAR (Summer of Learning And Research) program (to SAB).

## Supplementary Materials

Supplementary Figures S1 thru S13

Supplementary Table S1

Table legends

**Fig. S1.**
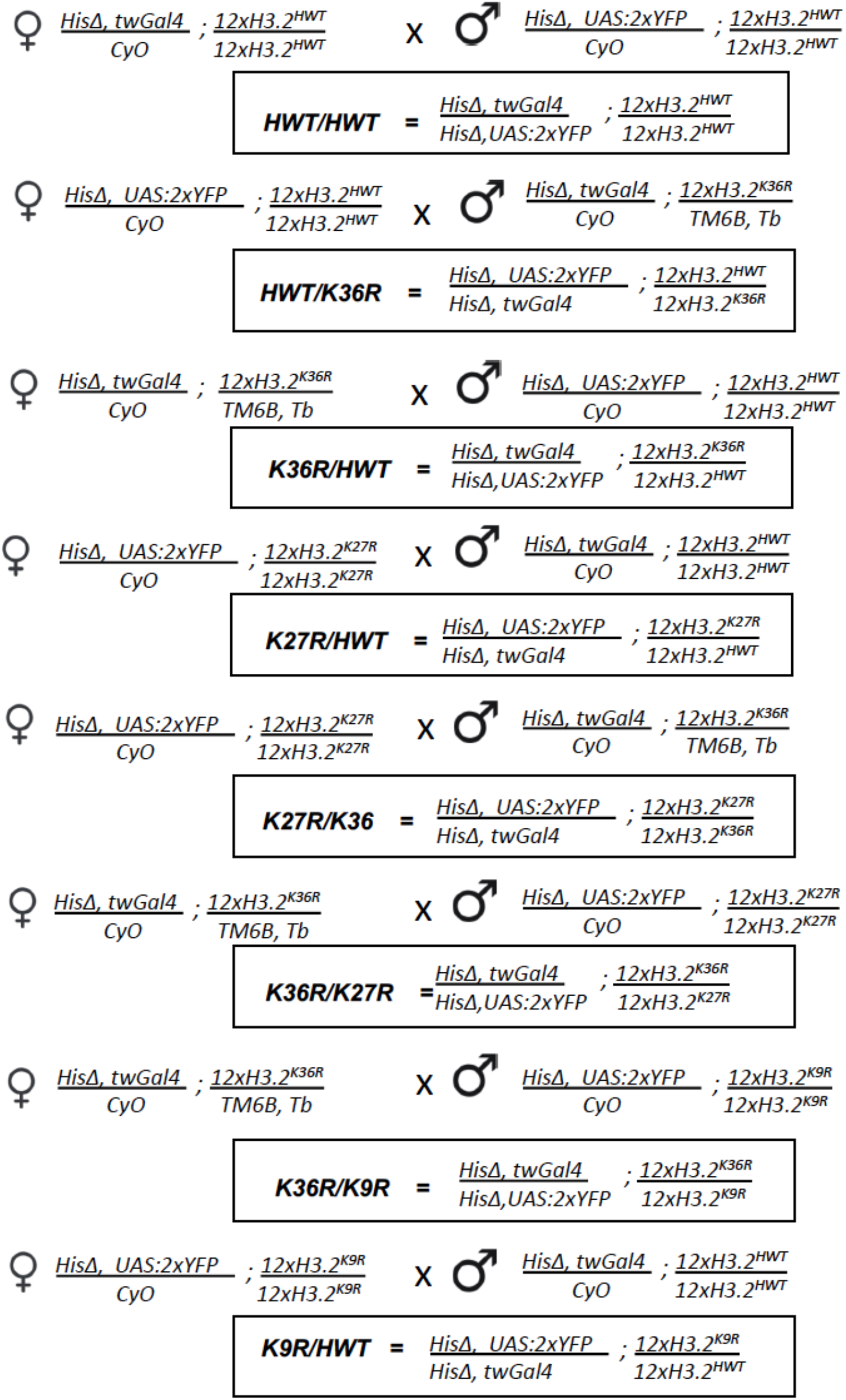
Crossing Schemes to Generate Genotypes for Figure 1. Crossing schemes used to generate the eight genotypes assayed in Figure 1B. Maternal and paternal genotypes are depicted above the boxes containing experimental genotypes. Experimental genotypes are indicated in both full and abbreviated forms.

**Fig. S2.**
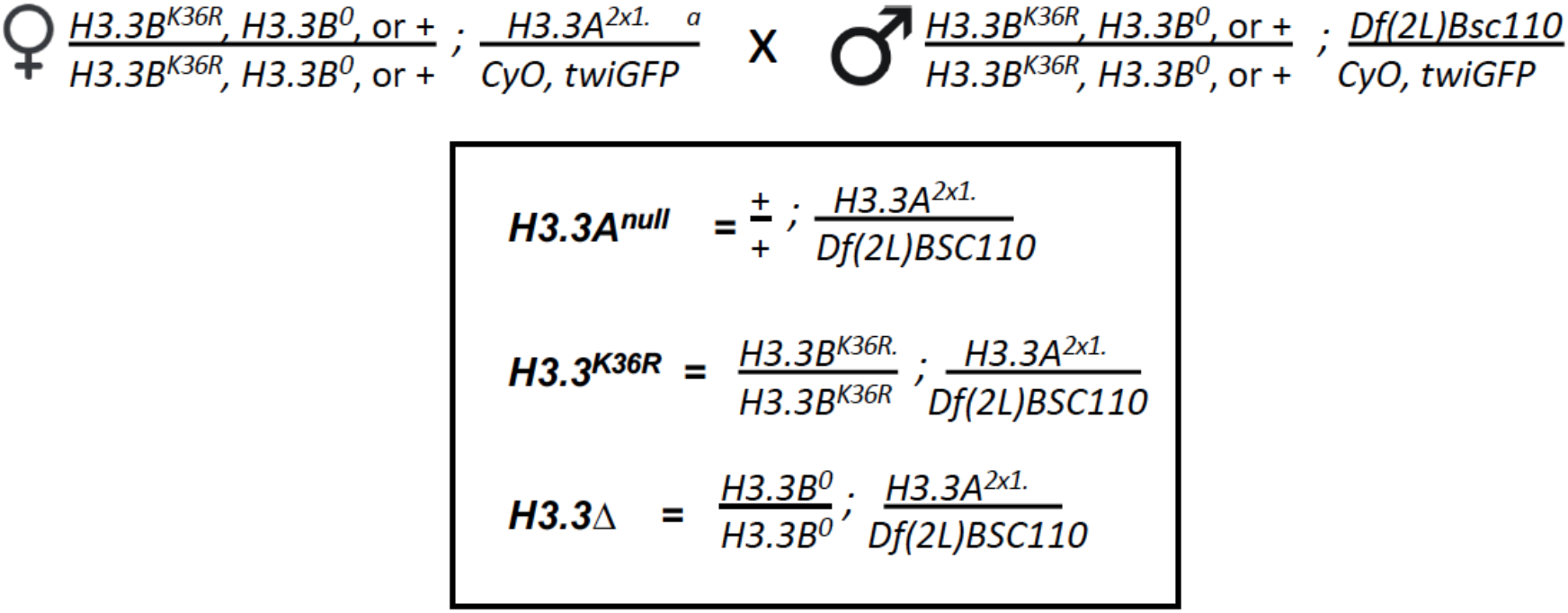
Crossing Schemes to Generate Genotypes for Figure 2. Crossing schemes used to generate 3 genotypes assayed in Figure 2A. Annotated as in Fig. S1. Note to obtain an *H3.3A*^*null*^ status for all, a deficiency (*Df(2L)BSC110*) was employed in trans to a null allele of *H3.3A*. Larvae were selected by GFP negative status.

**Fig. S3.**
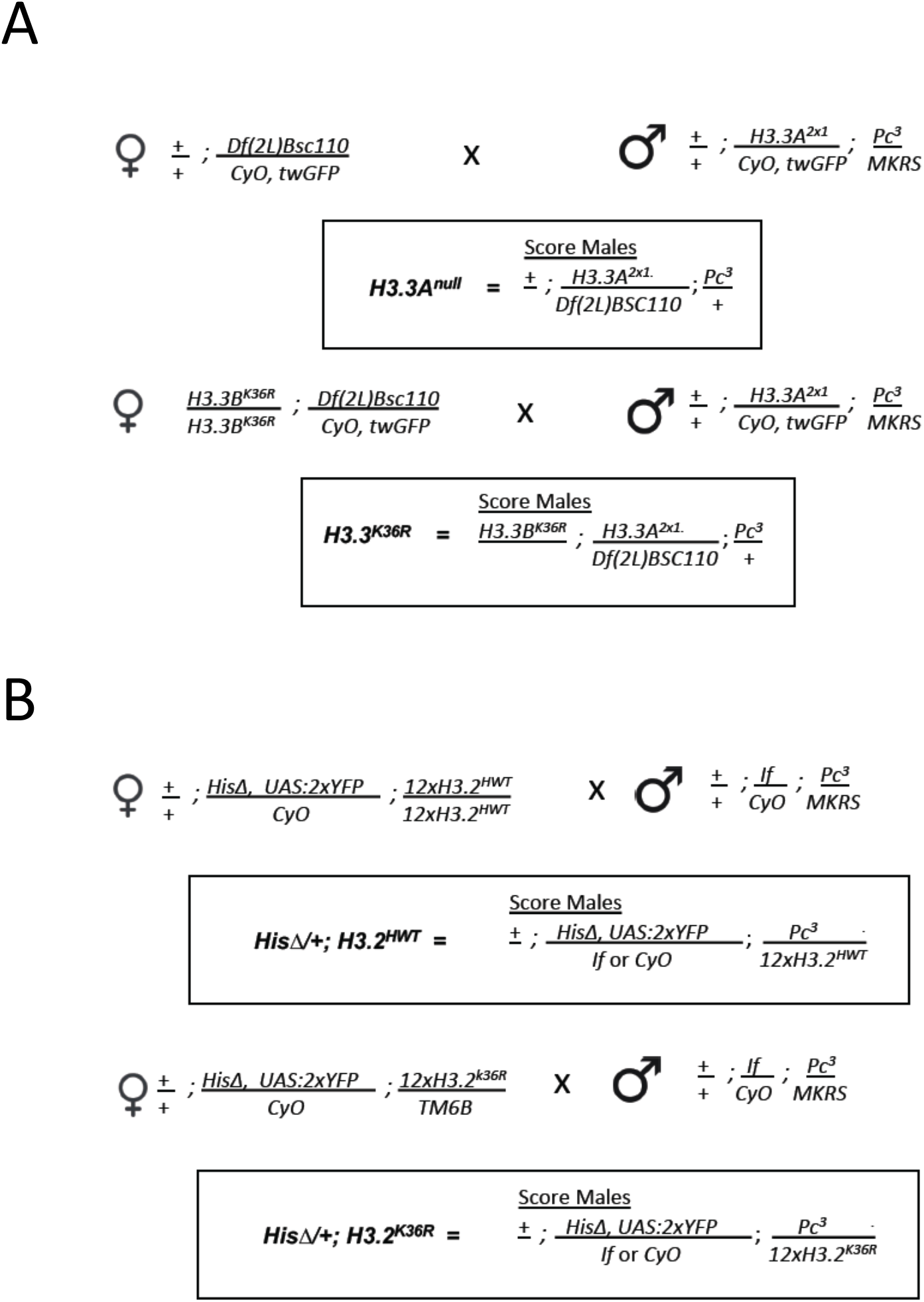
Crossing Schemes to Generate Genotypes for Figures 2, 3, and S5. Crossing schemes used to generate 4 genotypes assayed in Figures 2, 3, and S5. Annotated as in Fig. S1. **(A)** Crosses pertaining to *H3.3*^*K26R*^ mutant and control. Note that the *HisC* locus is wild type. Experimental progeny were Cy and GFP negative. **(B)** Crosses pertaining to *H3.2*^*K26R*^ mutant and control. Note that both *If* and *CyO* offspring were scored.

**Fig. S4.**
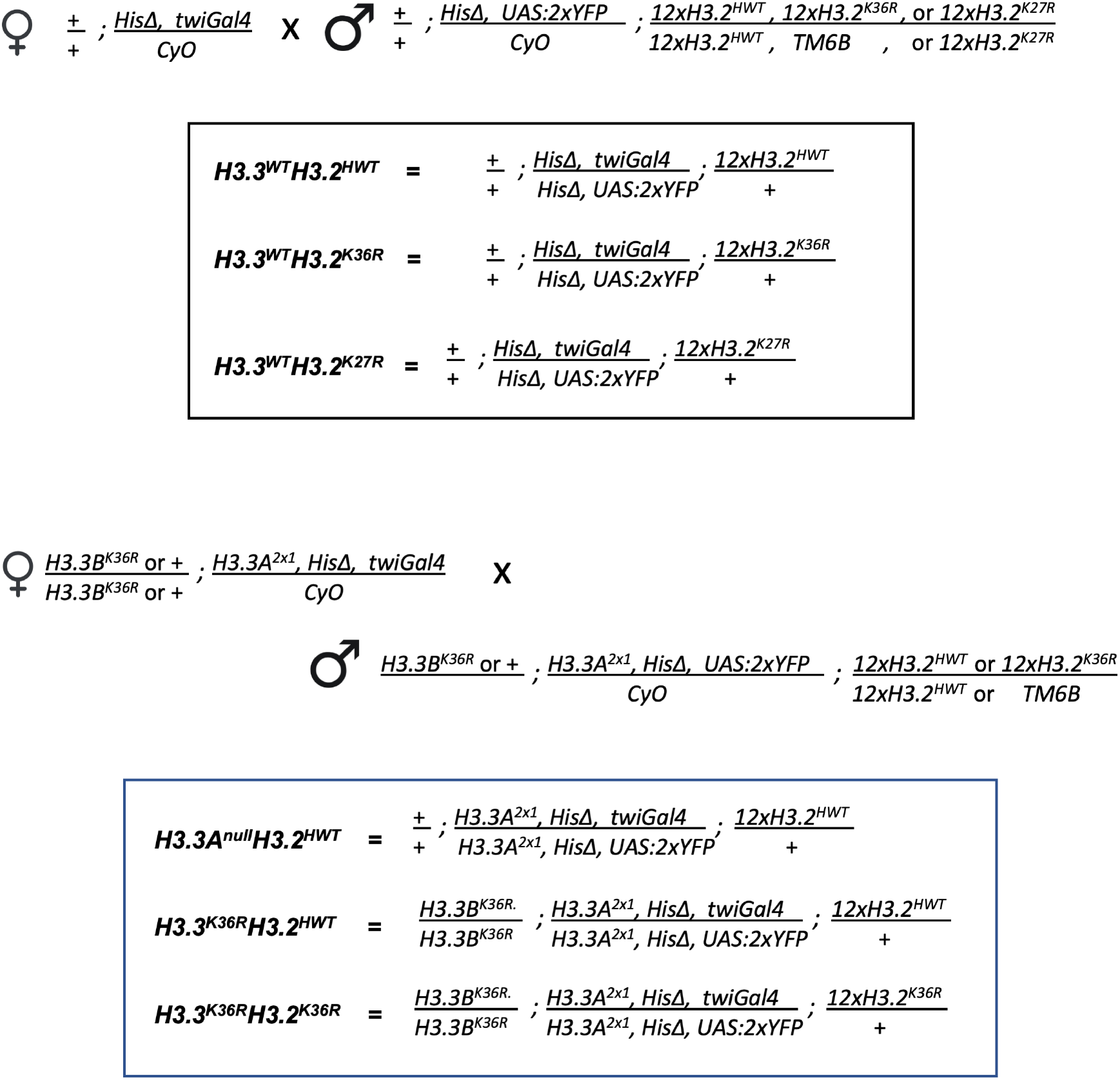
Crossing Schemes to Generate Genotypes for Figures 4-7. Crossing schemes used to generate 3 genotypes assayed in Figure 1B. Annotated as in Fig. S1. Note where applicable that to obtain an *H3.3A*^*null*^ status, the *H3.3A*^*2×1*^ allele is homozygous. All animals are in a *HisΔ* plus 12x histone replacement transgenic background. Larvae were selected by YFP positive status.

**Fig. S5.**
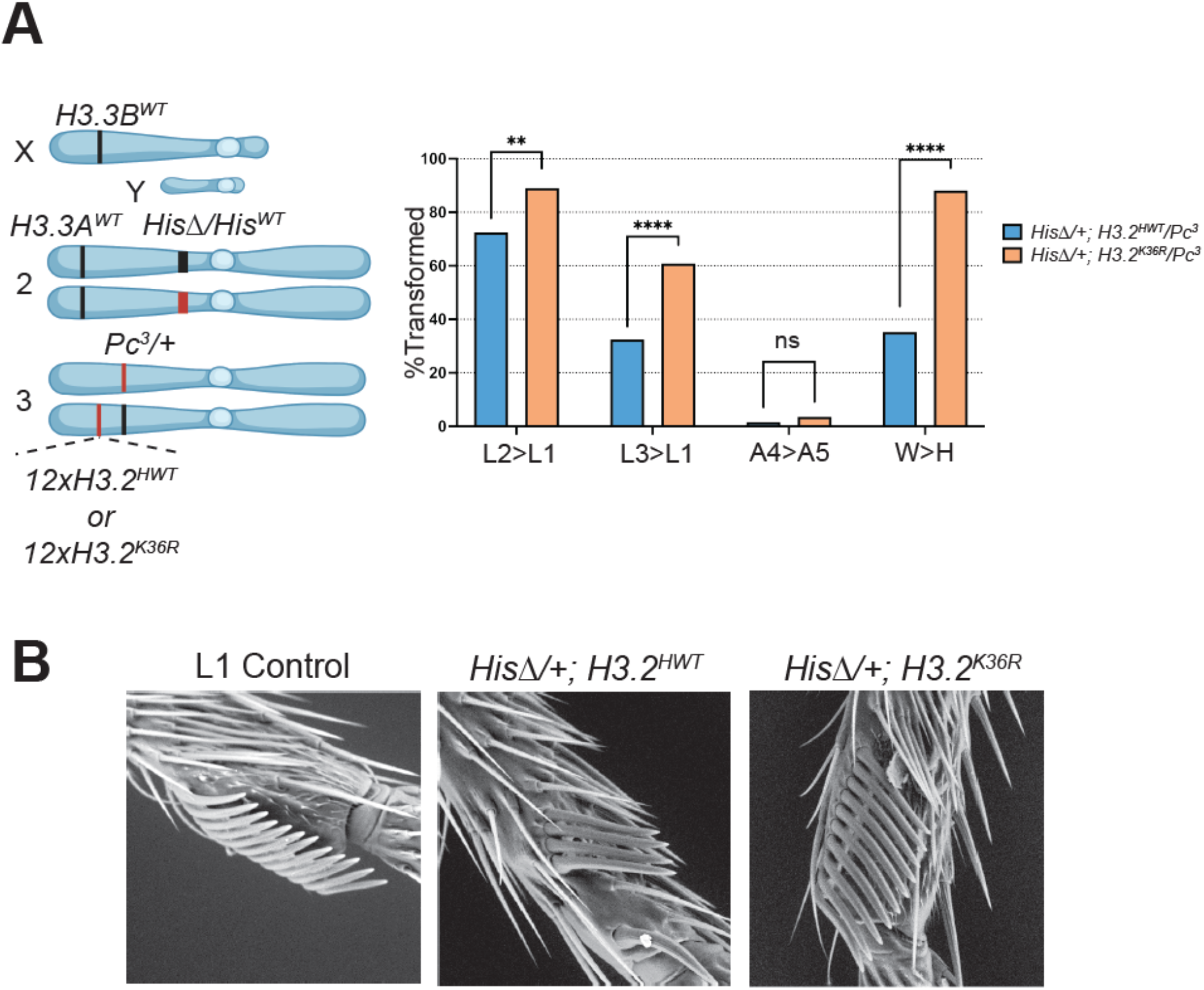
Ectopic expression of *H3.2*^*K36R*^ Genes Enhances Pc Transformations in Adult Flies. Either *H3.2*^*K36R*^ and control histone genotypes were combined with a heterozygous Pc^3^ mutation and scored for four characteristic PcG homeotic transformations. For full genetic scheme, see Fig. S2. T2-T1 (leg 2 to leg 1), T3-T1 (leg 3 to leg 4), A4-A5 (abdominal segment 4 to abdominal segment 5), and W-H (wing to haltere) transformations were scored for each genotype. Notably, for the *H3.2*^*K36R*^ analyses, the *HisC* locus was heterozygous to allow animals to reach adulthood, producing a ratio of *H3.2*^*WT*^ to *H3.2*^*K36R*^ genes of ∼10:1 (endogenous *HisC* locus contains ∼110 genes). **(A)** To the left, a summary of the genetic scheme for the *H3.2*^*K36R*^ and *Pc*^*3*^ genetic interaction experiment, created using BioRender.com. To the right, % Transformed for 4 homeotic transformations is plotted for each genotype. N value for number of flies scored for the *H3.2*^*K36R*^ genotype (n=55) and for the control (n=62). Note, for T2-T1 and T3-T1, each appendage was scored separately, effectively doubling the n value for these transformations. GraphPad Prism was used to calculate a χ^2^ value for each transformation. Significance is abbreviated as: *=p<0.05, **=p<0.01, ***=p<0.001, ****=p<0.0001. **(B)** Image of a typical T2-T1 transformation for each genotype collected by scanning electron microscopy at 250x magnification.

**Fig. S6.**
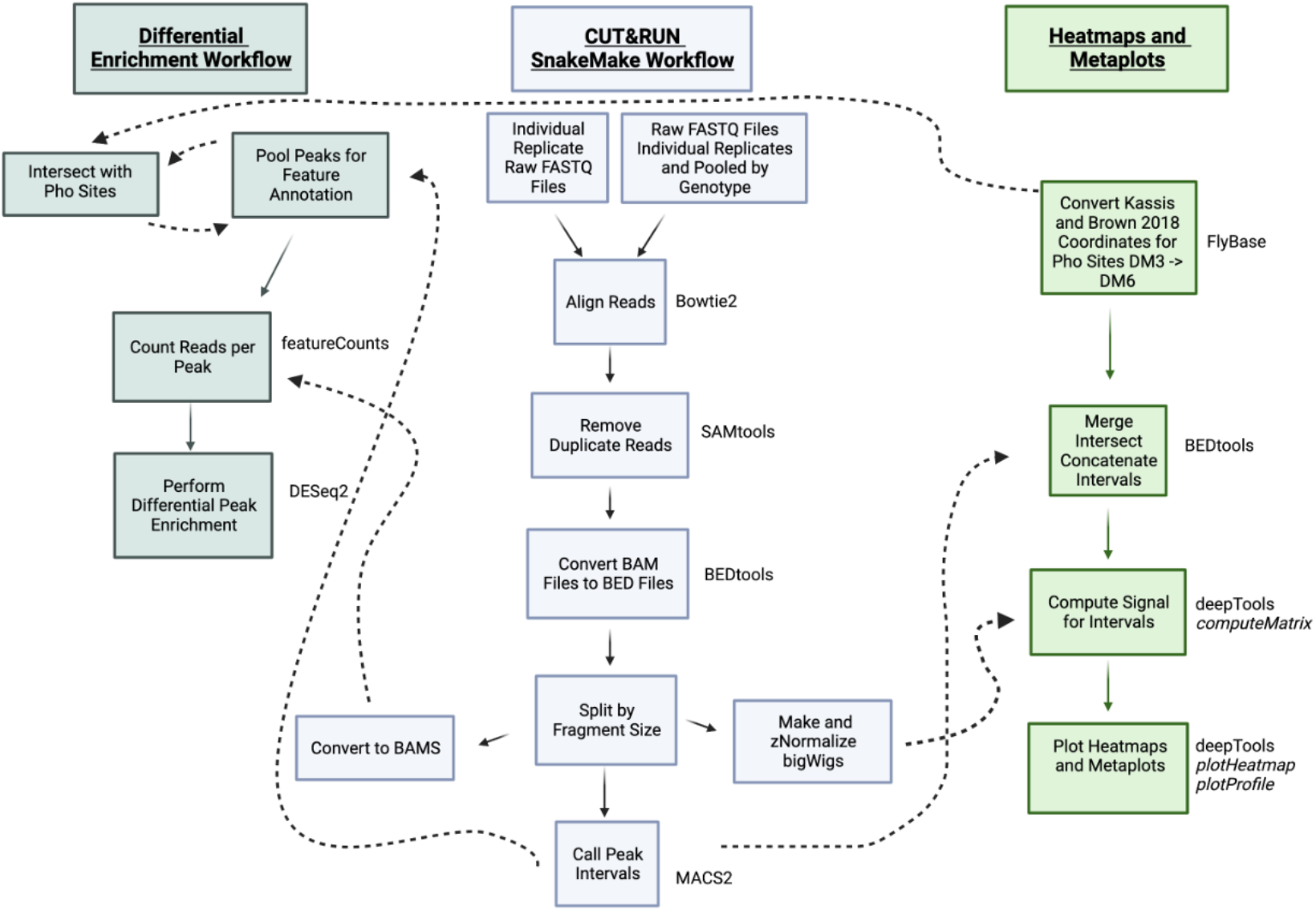
Bioinformatic Workflow for CUT&RUN Analysis. CUT&RUN data were processed as in https://github.com/mckaylabunc/cutNrun-pipeline with minor modifications. Briefly, QC was performed with FastQC (fastqc/0.11.7) (*110*) and FastQ Screen (v0.11.1) (*111*). Reads were aligned with Bowtie2 (v2.3.4.1) (*112*) to the DM6 reference genome. SAMtools (v1.10) was used to remove duplicate reads and sort BAM files (*113*). BAM files were converted to BED files using BEDTools (v2.26) (*109*). These BED files were separated by fragment length (20-120bp, short fragments; 150-700bp, long fragments) and converted to separate fragment size binned BAM files. Bedgraphs were generated using BEDTools (v2.26) (*109*) and wigToBigWig (*114*) was used to convert BED files to bigWigs. BigWig files were RPGC (reads per genome coverage) normalized, and further transformed by z-normalization. Peak calling was performed using MACS2 (v2.1.2) (*115*) without the use of an IgG or DNA control file. Differential peak analysis was performed using featureCounts (*106*) and DESeq2 (v1.34.0) (*107*). Details for generating intervals for each differential analysis (broad domains-Fig 4B, concatenated short fragment peaks from supernatant-Fig S11, short fragment peaks with Pho binding-Figs. 5C&S10B) can be found in Fig. S8. All genotypes were included to build each DESeq2 model. After models were created, differential analysis was performed between specified genotype comparisons of interest. For broad domains (Fig. 4B), pellet reads of all fragment sizes were used for DESeq2 analysis. For PRE based analyses in Fig 5C, S10B, and S11, only short fragment (20-120bp) pellet reads were analyzed with DESeq2. All heatmaps and metaplots were generated from pooled bigWigs for each genotype using the deepTools (v3.5.1) package (*108*) and the reference-point option, rather than scale-region. Details for reference-point selection can be found in Fig. S8. Details for which files and parameters were used to produce heatmaps and metaplots can be found in the *Bioinformatic Analyses* section of Materials and Methods. This figure was created using BioRender.com.

**Fig. S7.**
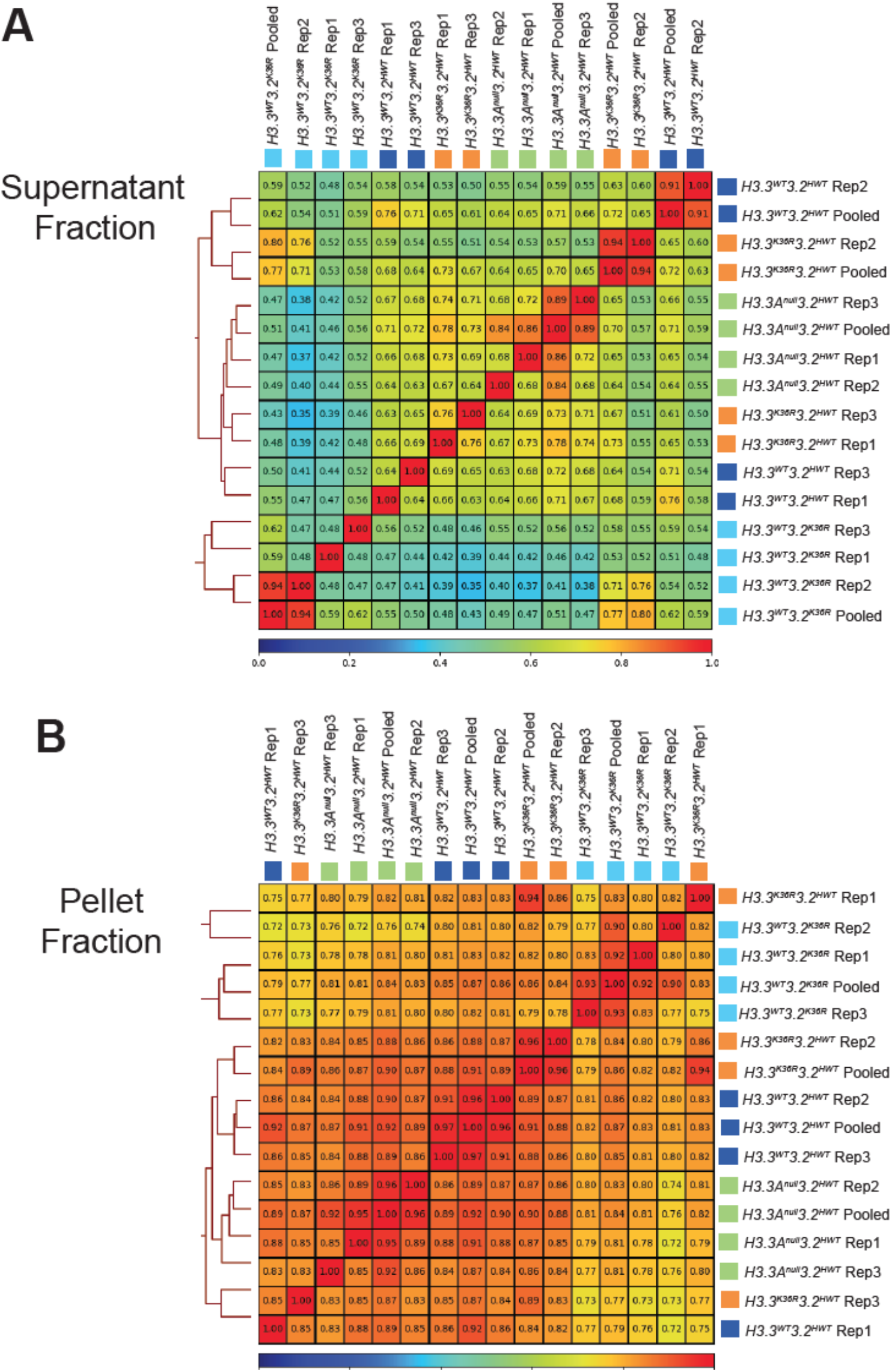
Spearman Correlations for CUT&RUN Replicates. The deepTools (v3.5.1) package (*108*), was used to calculate Spearman correlations between bigWig files from CUT&RUN replicates. **(A)** Supernatant fraction **(B)** Pellet fraction. Spearman correlations between replicates from the same genotype are higher from within the pellet fraction.

**Fig. S8.**
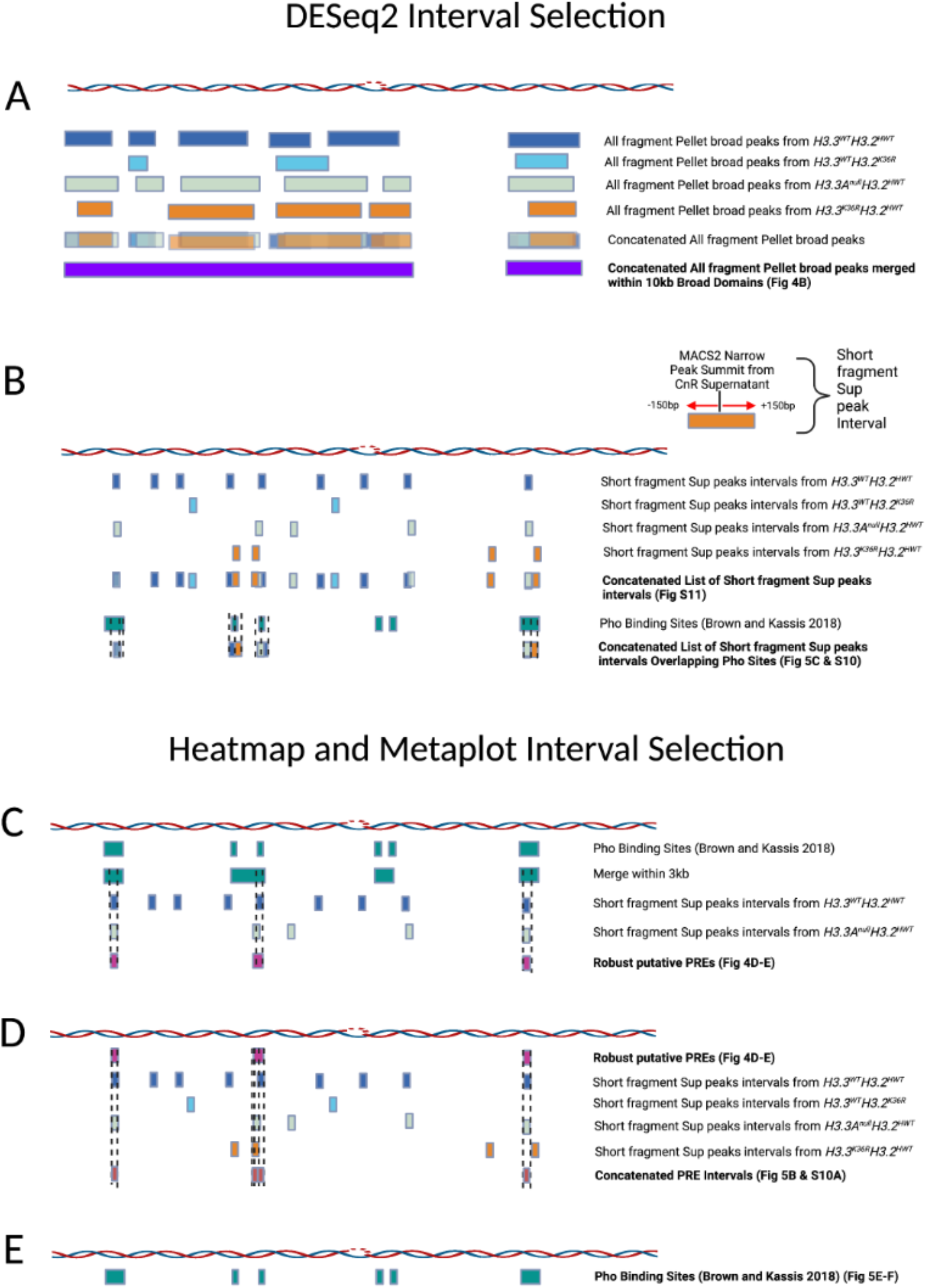
Interval Selection for Bioinformatic Analyses. BEDtools (v.2.3.0) (*109*) was used for intersecting and concatenating intervals from BED files generated from 1) Flybase Coordinate Converter Tool 2) MACS2 (*115*). **(A)** For DESeq2 analysis of broad domains (Fig. 4B), broad peaks output by MACS2 for each genotype were concatenated with Bedtoolsr (2.30.0-4) (*116*). Concatenated peaks were merged within 10kb to produce the broad domains used for the final analysis. **(B)** For narrow peak summit intervals in PRE analysis, short fragment supernatant narrow peak summits were extended +150bp and a master list concatenated from all four genotypes was used for the DESeq2 in analysis in Fig S11. For DESeq2 analyses of putative PRE regions with Pho binding capability, this list was intersected with Pho binding sites, and intervals overlapping Pho were analyzed in Fig 5C and S10B. **(C)** For heatmaps and metaplots in Fig4D-E, Pho binding regions from Brown and Kassis 2018 (*59*) were merged within 3kb with BEDTools (v.2.3.0). K27me3 MACS2 narrow peak summits + 150bp intervals from each control genotype that intersected these merged Pho intervals was compiled with BEDTools (v.2.3.0). Lists for both controls were intersected with BEDTools to generate a final list of “robust putative PREs” used for analysis in Fig. 4D. **(D)** For Figs. 5B & S10A metaplots, a master list of intervals from MACS2 narrow peak summits + 150bp for *H3.3*^*WT*^*H3.2*^*HWT*^, *H3.3*^*WT*^*H3.2*^*K36R*^, *H3.3A*^*null*^*H3.2*^*HWT*^, and *H3.3*^*K36R*^*H3.2*^*HWT*^ genotypes was concatenated. All intervals overlapping the “robust PREs” were used for subsequent metaplots. **(E)** For Figs. 5E-F, all unmerged Pho binding intervals (*59*) were sorted into 2 bed files by K27me3 status using the K27me3 broad domain peak annotation used for Fig. 4B with BEDTools (v.2.3.0). This figure was created using BioRender.com.

**Fig. S9.**
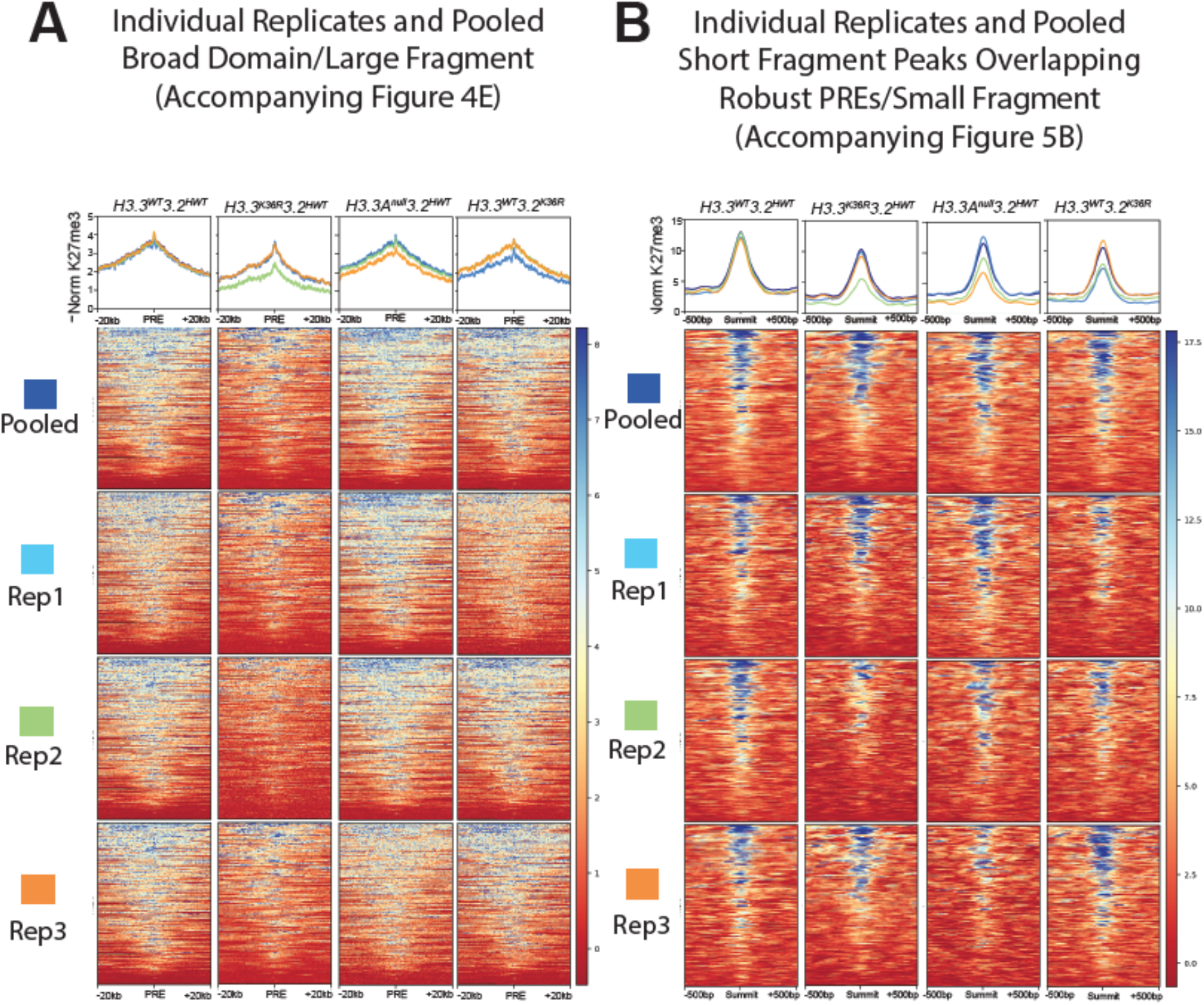
Metaplots and Heatmaps for Individual CUT&RUN Replicates. Analyses from Fig. 4E&5B were repeated with z-score normalized bigWigs from individual replicates alongside pooled bigWigs to assess the degree of variability between samples within the same genotype. **(A)** For broad domain analysis of large fragments, there is little variability between replicates, and pooled replicates overlay cleanly with the majority of individual replicates if one replicate is slightly lower. **(B)** Signal from short fragment pellet reads is generally more variable between replicates making precise quantification difficult.

**Fig. S10.**
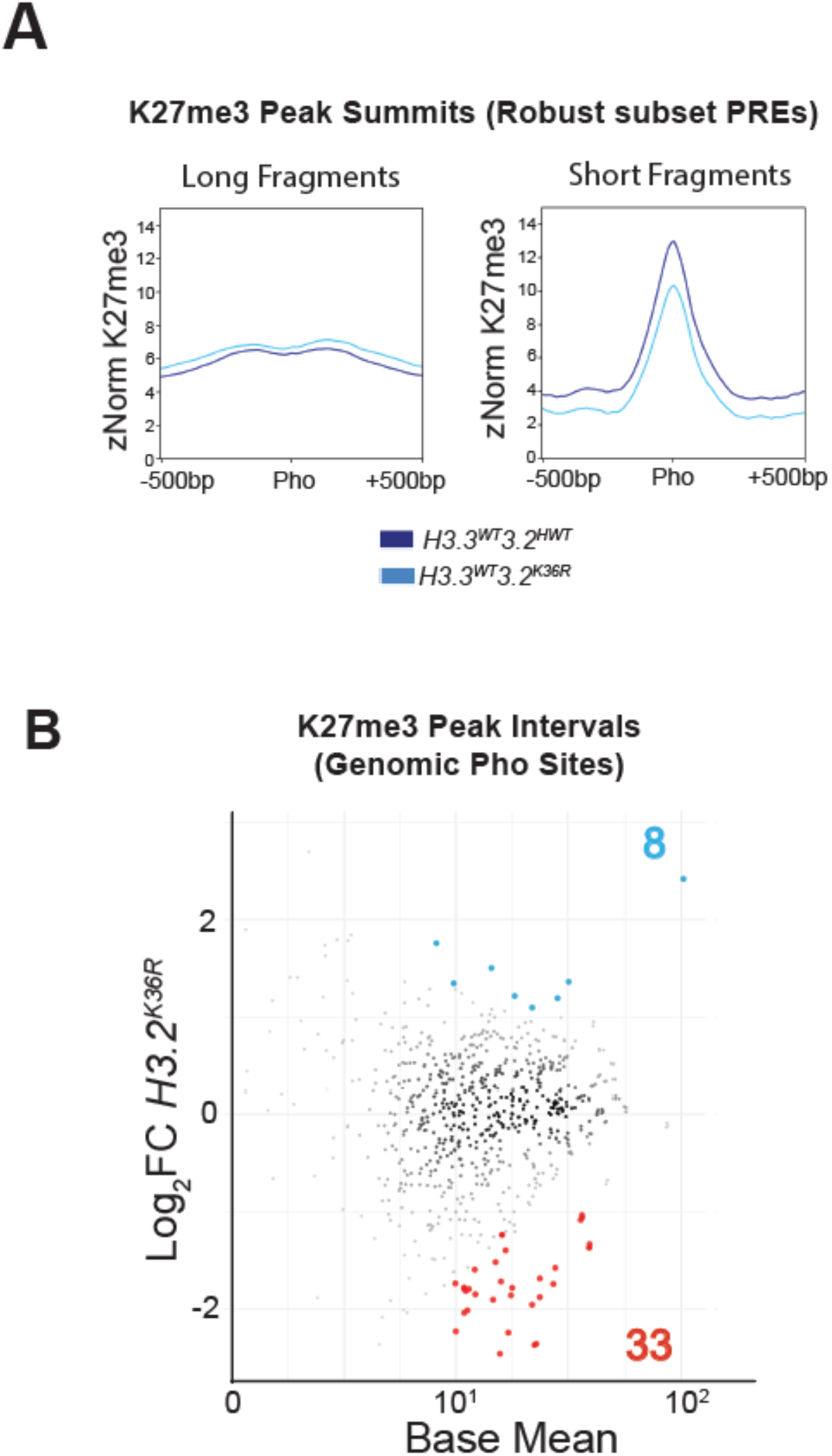
H3K27me3 Directed Cleavage is Unaltered at PREs in the *H3.2*^*K36R*^ Mutant. **(A)** Metaplots of K27me3 directed CUT&RUN signal + 500bp around peak summits called from all 4 genotypes that overlap robust PREs intervals identified in Figure 4 (n=344), see Fig. S8 for details. Separate plots were generated for long and short fragments. The *H3.3*^*WT*^*H3.2*^*K36R*^ mutant is plotted alongside the *H3.3*^*WT*^*H3.2*^*HWT*^ control. **(B)** DESeq2 analysis for peak summit intervals identified from all 4 genotypes overlapping Kassis Pho binding sites genome-wide (n=711) for the *H3.3*^*K36R*^*H3.2*^*HWT*^ mutant vs. control, annotated as in Fig 4B.

**Fig. S11.**
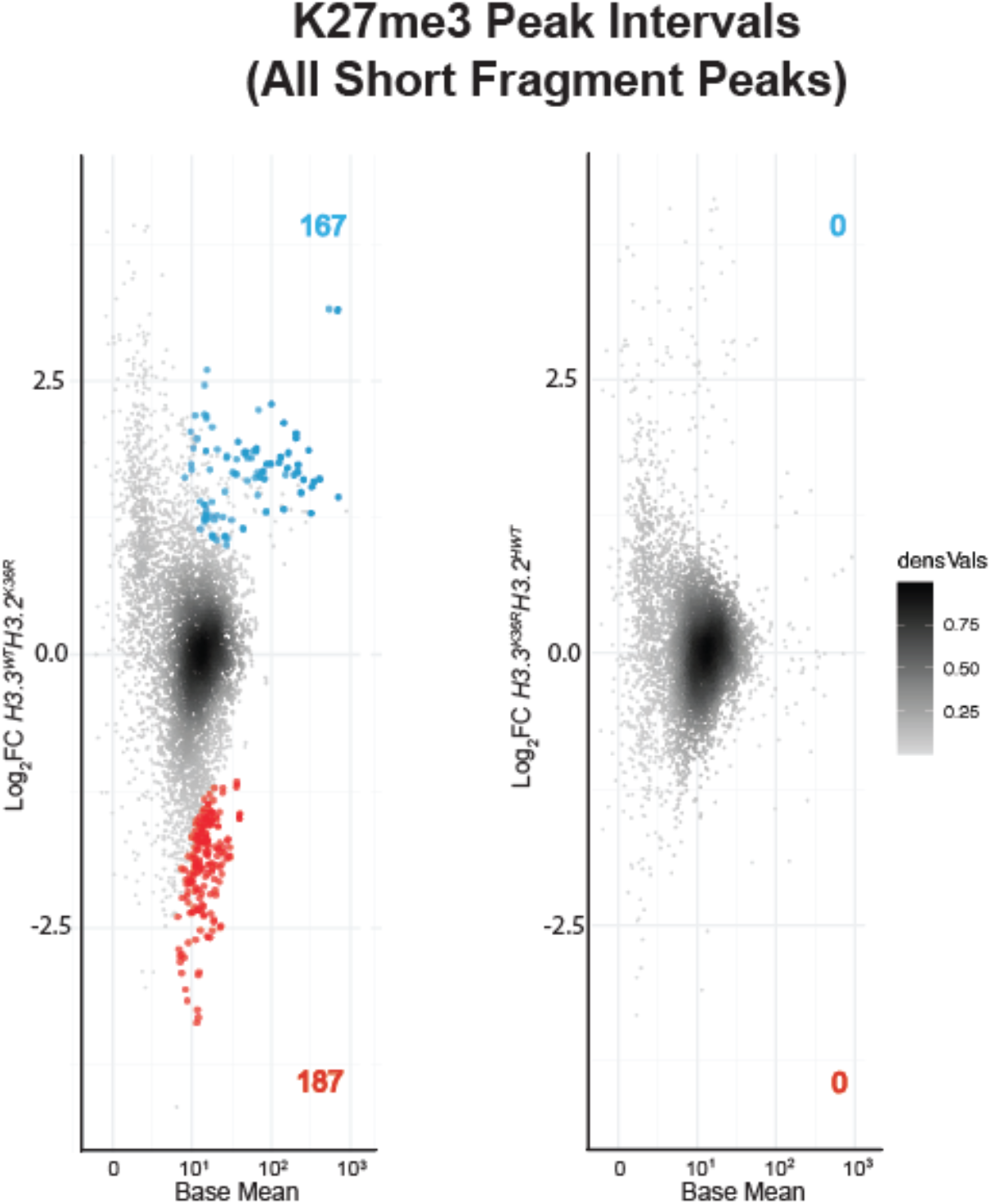
Images Used for Staging Embryos in Figure 3. **(A)** Stage 16 embryos of H3K36R mutant genotypes and controls were fixed and stained with anti-AbdB antibodies. Embryos were stained with anti-GFP antibodies to detect YFP for staging and genotype selection. For each embryo, a single slice from the anti-GFP channel was used for staging the embryos in Figure 3A. Scale bar = 50µm. **(B)** Same as in A, except stained with anti-Ubx instead of anti-AbdB, and from the embryos depicted in Figure 3B.

**Fig. S12.**
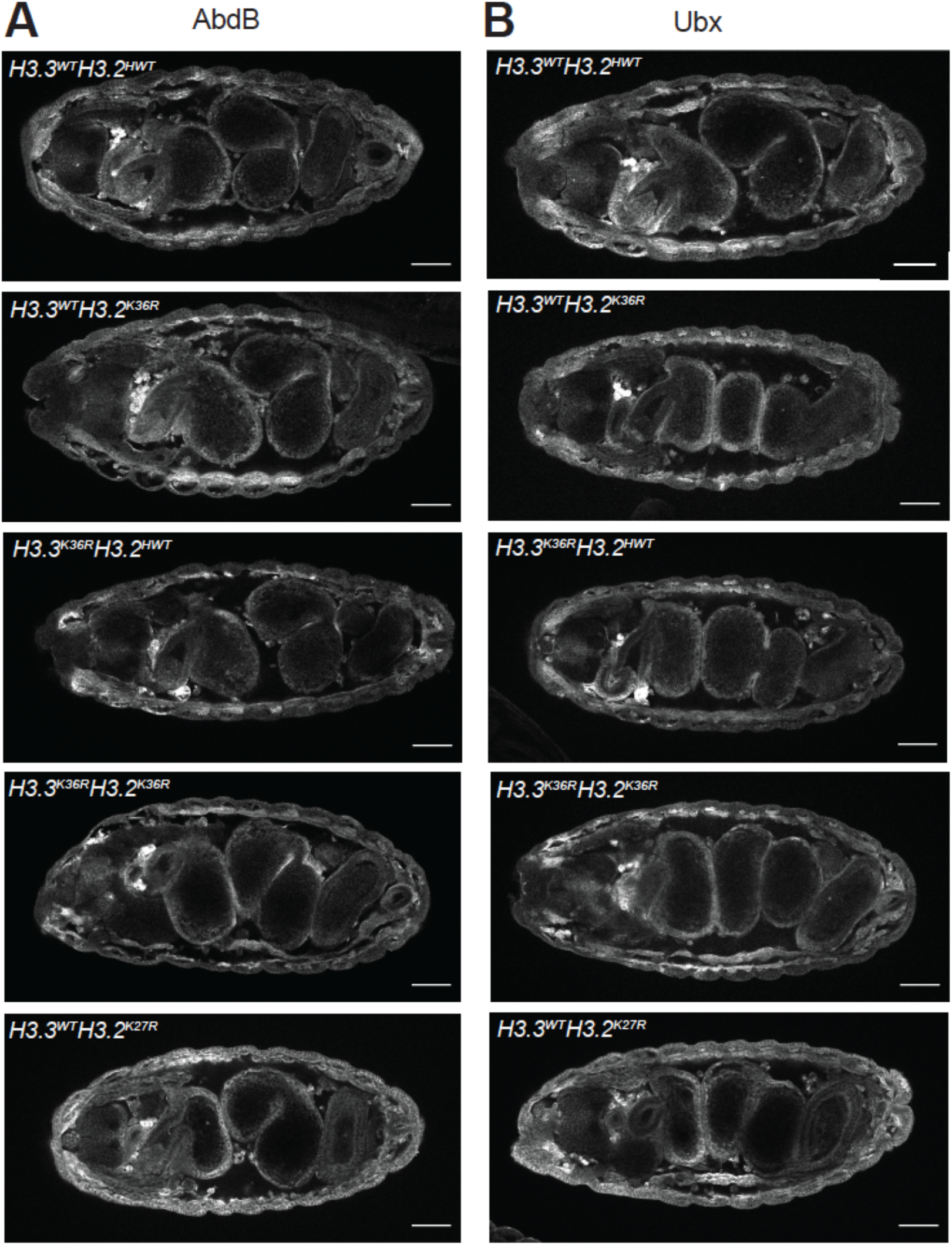
Embryos with Combined Mutation of *H3.3*^*K36R*^ and *H3.2*^*K36R*^ Exhibit Moderate Synergistic Derepression of ANTP. Stage 16 embryos of H3K36R mutant genotypes and controls were fixed and stained with anti-ANTP antibodies. Embryos were stained with anti-GFP antibodies to detect YFP for staging and genotype selection. Scale bar = 50µm. **(A)** Representative Anti-ANTP staining for 5 genotypes. Brackets indicate the expected boundary of ANTP expression in wild type embryos. Filled arrows highlight individual cells exhibiting anterior derepression of ANTP. Black arrows indicate derepressed cells. Overall, individual H3.2 and H3.3 mutants closely resemble *H3.3*^*WT*^*H3.2*^*HWT*^ negative controls, while *H3.3*^*K36R*^*H3.2*^*K36R*^ were generally intermediate between *H3.3*^*WT*^*H3.2*^*HWT*^ and *H3.3*^*WT*^*H3.2*^*K27R*^ controls. **(B)** Single slice from anti-GFP channel staining YFP for same embryos in A depicting staging.

**Fig. S13.**
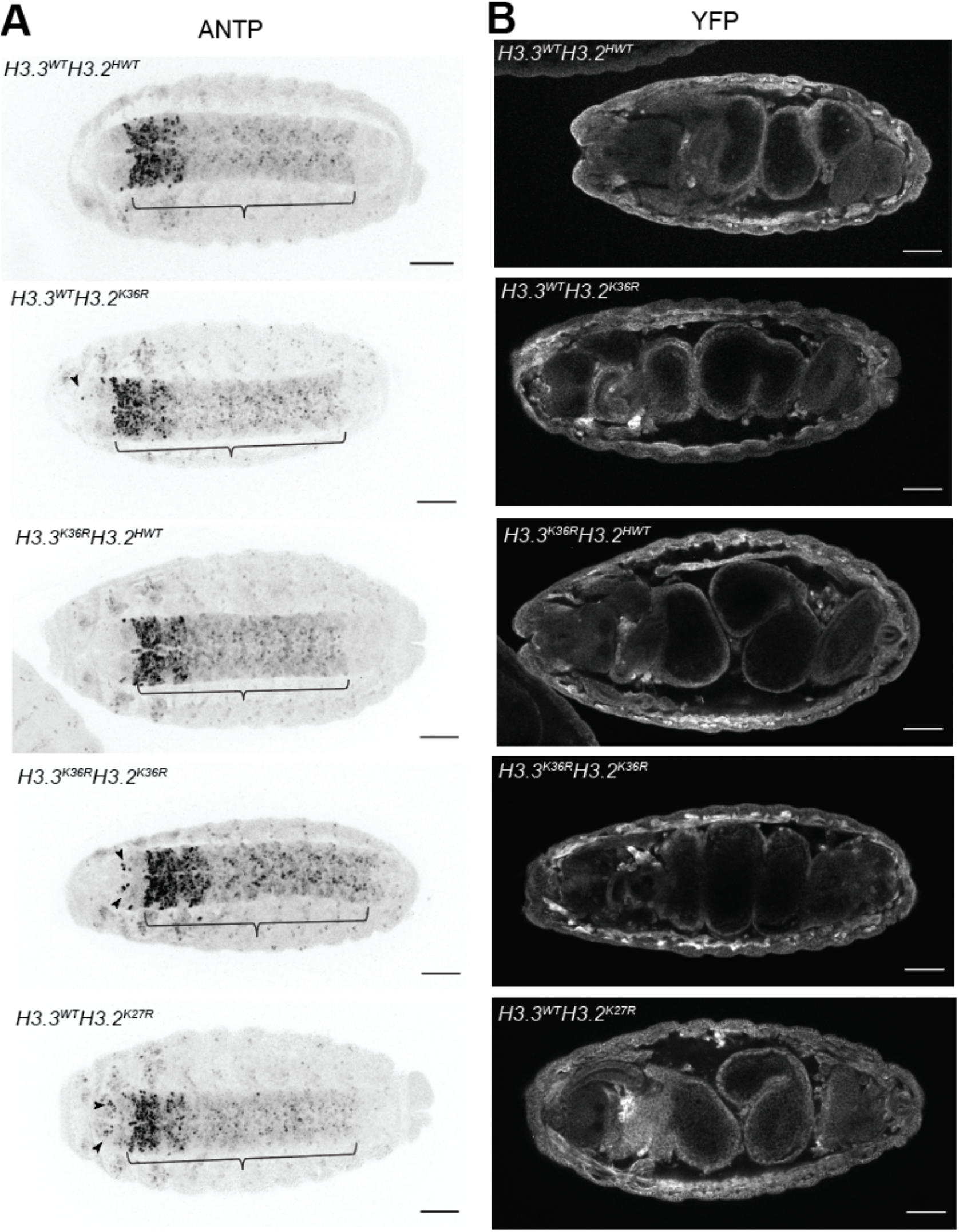
Embryos with Combined Mutation of *H3.3*^*K36R*^ and *H3.2*^*K36R*^ Exhibit Moderate Synergistic Derepression of ANTP. Stage 16 embryos of H3K36R mutant genotypes and controls were fixed and stained with anti-ANTP antibodies. Embryos were stained with anti-GFP antibodies to detect YFP for staging and genotype selection. Scale bar = 50µm. **(A)** Representative Anti-ANTP staining for 5 genotypes. Brackets indicate the expected boundary of ANTP expression in wild type embryos. Filled arrows highlight individual cells exhibiting anterior derepression of ANTP. Black arrows indicate derepressed cells. Overall, individual H3.2 and H3.3 mutants closely resemble *H3.3*^*WT*^*H3.2*^*HWT*^ negative controls, while *H3.3*^*K36R*^*H3.2*^*K36R*^ were generally intermediate between *H3.3*^*WT*^*H3.2*^*HWT*^ and *H3.3*^*WT*^*H3.2*^*K27R*^ controls. **(B)** Single slice from anti-GFP channel staining YFP for same embryos in A depicting staging.

**Table S1.** Shorthand nomenclature for genotypes used in this study. A key is provided here for easy reference. The left column indicates the shorthand genotype used in the text and figures. The right column lists the full genotype with all relevant alleles and transgenes.

| Nomenclature | Genotype |
| --- | --- |
| For Figure 1B |  |
| *HWT/HWT* | $\frac{His\Delta, twiGal4}{His\Delta, UAS:YFP}; \frac{12xH3.2^{HWT}}{12xH3.2^{HWT}}$ |
| *HWT/K36R* | $\frac{His\Delta, twiGal4}{His\Delta, UAS:YFP}; \frac{12xH3.2^{HWT}}{12xH3.2^{K36R}}$ |
| *K36R/HWT* | $\frac{His\Delta, twiGal4}{His\Delta, UAS:YFP}; \frac{12xH3.2^{K36R}}{12xH3.2^{HWT}}$ |
| *K36R/K27R* | $\frac{His\Delta, twiGal4}{His\Delta, UAS:YFP}; \frac{12xH3.2^{K36R}}{12xH3.2^{K27R}}$ |
| *K36R/K9R* | $\frac{His\Delta, twiGal4}{His\Delta, UAS:YFP}; \frac{12xH3.2^{K36R}}{12xH3.2^{K9R}}$ |
| *K27R/K36R* | $\frac{His\Delta, twiGal4}{His\Delta, UAS:YFP}; \frac{12xH3.2^{K27R}}{12xH3.2^{K36R}}$ |
| *K27R/HWT* | $\frac{His\Delta, twiGal4}{His\Delta, UAS:YFP}; \frac{12xH3.2^{K36R}}{12xH3.2^{HWT}}$ |
| *K9R/HWT* | $\frac{His\Delta, twiGal4}{His\Delta, UAS:YFP}; \frac{12xH3.2^{K9R}}{12xH3.2^{HWT}}$ |
| For Figures 1C, and 2-6 |  |
| *H3.3A<sup>null</sup>* | $\frac{+}{+}; \frac{H3.3A^{2x1}}{Df(2L)Bsc110}; \frac{+}{+}$ |
| *H3.3<sup>K36R</sup>* | $\frac{H3.3B^{K36R}}{H3.3B^{K36R}}; \frac{H3.3A^{2x1}}{Df(2L)Bsc110}; \frac{+}{+}$ |
| *H3.3Δ* | $\frac{H3.3B^0}{H3.3B^0}; \frac{H3.3A^{2x1}}{Df(2L)Bsc110}; \frac{+}{+}$ |
| *H3.3<sup>WT</sup>H3.2<sup>HWT</sup>* | $\frac{+}{+}; \frac{His\Delta, twiGal4}{His\Delta, UAS:YFP}; \frac{12xH3.2^{HWT}}{+}$ |
| *H3.3<sup>WT</sup>H3.2<sup>K36R</sup>* | $\frac{+}{+}; \frac{His\Delta, twiGal4}{His\Delta, UAS:YFP}; \frac{12xH3.2^{K36R}}{+}$ |
| *H3.3<sup>null</sup>H3.2<sup>HWT</sup>* | $\frac{+}{+}; \frac{H3.3A^{2x1}, His\Delta, twiGal4}{H3.3A^{2x1}, His\Delta, UAS:YFP}; \frac{12xH3.2^{HWT}}{+}$ |
| *H3.3<sup>K36R</sup>H3.2<sup>HWT</sup>* | $\frac{H3.3B^{K36R}}{H3.3B^{K36R}}; \frac{H3.3A^{2x1}, His\Delta, twiGal4}{H3.3A^{2x1}, His\Delta, UAS:YFP}; \frac{12xH3.2^{HWT}}{+}$ |
| *H3.3<sup>K36R</sup>H3.2<sup>K36R</sup>* | $\frac{H3.3B^{K36R}}{H3.3B^{K36R}}; \frac{H3.3A^{2x1}, His\Delta, twiGal4}{H3.3A^{2x1}, His\Delta, UAS:YFP}; \frac{12xH3.2^{K36R}}{+}$ |
| *H3.3<sup>WT</sup>H3.2<sup>K27R</sup>* | $\frac{+}{+}; \frac{His\Delta, twiGal4}{His\Delta, UAS:YFP}; \frac{12xH3.2^{K27R}}{+}$ |

**Table S2. DESeq2 Output for K27me3 CUT&RUN for Broad Domains**. DESeq2 output accompanying Fig. 4B. Differential peak analysis by DESeq2 on broad H3K27me3 domains (details in Fig.S8) using all-fragment BAM files from the pellet fraction. Separate columns comparing *H3.3*^*WT*^*H3.2*^*K36R*^ vs. *H3.3*^*WT*^*H3.2*^*HWT*^ control and *H3.3*^*K36R*^*H3.2*^*HWT*^ vs. *H3.3A*^*null*^*H3.2*^*HWT*^ are included.

**Table S3. DESeq2 Output for K27me3 CUT&RUN for Short Fragment Peak Intervals Overlapping Pho**. DESeq2 output accompanying Fig. 5C and S10B. Differential peak analysis by DESeq2 on short fragment peak intervals from H3K27me3 domains CUT&RUN that overlap Pho binding sites (details in Fig.S8) using small fragment BAM files from the pellet fraction. Separate columns comparing *H3.3*^*WT*^*H3.2*^*K36R*^ vs. *H3.3*^*WT*^*H3.2*^*HWT*^ control and *H3.3*^*K36R*^*H3.2*^*HWT*^ vs. *H3.3A*^*null*^*H3.2*^*HWT*^ are included.

**Table S4. DESeq2 Output for All K27me3 CUT&RUN for Short Fragment Peak Intervals**. DESeq2 output accompanying Fig. S11. Differential peak analysis by DESeq2 on short fragment peak intervals from H3K27me3 domains CUT&RUN (details in Fig.S8) using small fragment BAM files from the pellet fraction. Separate columns comparing *H3.3*^*WT*^*H3.2*^*K36R*^ vs. *H3.3*^*WT*^*H3.2*^*HWT*^ control and *H3.3*^*K36R*^*H3.2*^*HWT*^ vs. *H3.3A*^*null*^*H3.2*^*HWT*^ are included.

## References

1. N. Camacho-Ordonez, E. Ballestar, H. T. M. Timmers, B. Grimbacher, What can clinical immunology learn from inborn errors of epigenetic regulators? J Allergy Clin Immunol 147, 1602–1618 (2021).

2. E. Conway, E. Healy, A. P. Bracken, PRC2 mediated H3K27 methylations in cellular identity and cancer. Curr Opin Cell Biol 37, 42–48 (2015).

3. J. H. Lee, J.-H. Kim, S. Kim, K. S. Cho, S. B. Lee, Chromatin Changes Associated with Neuronal Maintenance and Their Pharmacological Application. Current Neuropharmacology 16, (2018).

4. A. Piunti, A. Shilatifard, The roles of Polycomb repressive complexes in mammalian development and cancer. Nature Reviews Molecular Cell Biology 22, 326–345 (2021).

5. A. Elsherbiny, G. Dobreva, Epigenetic memory of cell fate commitment. Curr Opin Cell Biol 69, 80–87 (2021).

6. J.-Y. Hwang, K. A. Aromolaran, R. S. Zukin, The emerging field of epigenetics in neurodegeneration and neuroprotection. Nature Reviews Neuroscience 18, 347–361 (2017).

7. K. R. Stewart-Morgan, N. Petryk, A. Groth, Chromatin replication and epigenetic cell memory. Nature Cell Biology 22, 361–371 (2020).

8. B. D. Strahl, C. D. Allis, The language of covalent histone modifications. Nature 403, 41–45 (2000).

9. D. J. McKay et al., Interrogating the function of metazoan histones using engineered gene clusters. Dev Cell 32, 373–386 (2015).

10. M. P. Meers et al., An Animal Model for Genetic Analysis of Multi-Gene Families: Cloning and Transgenesis of Large Tandemly Repeated Histone Gene Clusters. Methods Mol Biol 1832, 309–325 (2018).

11. R. L. Armstrong et al., Chromatin conformation and transcriptional activity are permissive regulators of DNA replication initiation in Drosophila. Genome Res 28, 1688–1700 (2018).

12. M. P. Meers et al., Histone gene replacement reveals a post-transcriptional role for H3K36 in maintaining metazoan transcriptome fidelity. Elife 6, (2017).

13. T. J. Penke, D. J. McKay, B. D. Strahl, A. G. Matera, R. J. Duronio, Direct interrogation of the role of H3K9 in metazoan heterochromatin function. Genes Dev 30, 1866–1880 (2016).

14. T. J. R. Penke, D. J. McKay, B. D. Strahl, A. G. Matera, R. J. Duronio, Functional Redundancy of Variant and Canonical Histone H3 Lysine 9 Modification in Drosophila. Genetics 208, 229–244 (2018).

15. B. Schuettengruber, H. M. Bourbon, L. Di Croce, G. Cavalli, Genome Regulation by Polycomb and Trithorax: 70 Years and Counting. Cell 171, 34–57 (2017).

16. J. A. Simon, R. E. Kingston, Occupying chromatin: Polycomb mechanisms for getting to genomic targets, stopping transcriptional traffic, and staying put. Mol Cell 49, 808–824 (2013).

17. R. T. Coleman, G. Struhl, Causal role for inheritance of H3K27me3 in maintaining the OFF state of a Drosophila HOX gene. Science 356, (2017).

18. A. R. Pengelly, Ö. Copur, H. Jäckle, A. Herzig, J. Müller, A histone mutant reproduces the phenotype caused by loss of histone-modifying factor Polycomb. Science 339, 698–699 (2013).

19. K. H. Hansen et al., A model for transmission of the H3K27me3 epigenetic mark. Nature Cell Biology 10, 1291–1300 (2008).

20. L. Jiao, X. Liu, Structural basis of histone H3K27 trimethylation by an active polycomb repressive complex 2. Science 350, aac4383–aac4383 (2015).

21. R. Margueron et al., Role of the polycomb protein EED in the propagation of repressive histone marks. Nature 461, 762–767 (2009).

22. M. Uckelmann, C. Davidovich, Not just a writer: PRC2 as a chromatin reader. Biochem Soc Trans 49, 1159–1170 (2021).

23. T. Zhang, S. Cooper, N. Brockdorff, The interplay of histone modifications - writers that read. EMBO Rep 16, 1467–1481 (2015).

24. N. P. Blackledge, R. J. Klose, The molecular principles of gene regulation by Polycomb repressive complexes. Nat Rev Mol Cell Biol 22, 815–833 (2021).

25. F. W. Schmitges et al., Histone methylation by PRC2 is inhibited by active chromatin marks. Mol Cell 42, 330–341 (2011).

26. W. Yuan et al., H3K36 methylation antagonizes PRC2-mediated H3K27 methylation. J Biol Chem 286, 7983–7989 (2011).

27. K. Finogenova et al., Structural basis for PRC2 decoding of active histone methylation marks H3K36me2/3. Elife 9, (2020).

28. E. Dorafshan et al., Ash1 counteracts Polycomb repression independent of histone H3 lysine 36 methylation. EMBO Rep, (2019).

29. S. J. Elsaesser, A. D. Goldberg, C. D. Allis, New functions for an old variant: no substitute for histone H3.3. Curr Opin Genet Dev 20, 110–117 (2010).

30. B. Loppin, F. Berger, Histone Variants: The Nexus of Developmental Decisions and Epigenetic Memory. Annu Rev Genet 54, 121–149 (2020).

31. A. Sakai, B. E. Schwartz, S. Goldstein, K. Ahmad, Transcriptional and developmental functions of the H3.3 histone variant in Drosophila. Curr Biol 19, 1816–1820 (2009).

32. J. A. Kassis, J. L. Brown. (Elsevier, 2013), pp. 83–118.

33. J. Muller, J. A. Kassis, Polycomb response elements and targeting of Polycomb group proteins in Drosophila. Curr Opin Genet Dev 16, 476–484 (2006).

34. G. Streubel et al., The H3K36me2 Methyltransferase Nsd1 Demarcates PRC2-Mediated H3K27me2 and H3K27me3 Domains in Embryonic Stem Cells. Mol Cell 70, 371–379 e375 (2018).

35. Y. Zheng et al., Total kinetic analysis reveals how combinatorial methylation patterns are established on lysines 27 and 36 of histone H3. Proc Natl Acad Sci U S A 109, 13549–13554 (2012).

36. P. B. Talbert, S. Henikoff, Histone variants on the move: substrates for chromatin dynamics. Nat Rev Mol Cell Biol 18, 115–126 (2017).

37. A. S. Akhmanova et al., Structure and expression of histone H3.3 genes in Drosophilamelanogaster and Drosophila-hydei. Genome 38, 586–600 (1995).

38. M. Hodl, K. Basler, Transcription in the absence of histone H3.3. Curr Biol 19, 1221–1226 (2009).

39. M. Leatham-Jensen et al., Lysine 27 of replication-independent histone H3.3 is required for Polycomb target gene silencing but not for gene activation. PLoS Genet 15, e1007932 (2019).

40. S. E. Celniker, S. Sharma, D. J. Keelan, E. B. Lewis, The molecular genetics of the bithorax complex of Drosophila: cis-regulation in the Abdominal-B domain. The EMBO journal 9, 4277–4286 (1990).

41. N. Javeed, N. J. Tardi, M. Maher, S. Singari, K. A. Edwards, Controlled expression of Drosophila homeobox loci using theHostile takeoversystem. Developmental Dynamics 244, 808–825 (2015).

42. U. Fresán, M. A. Rodríguez-Sánchez, O. Reina, V. G. Corces, M. L. Espinàs, Haspin kinase modulates nuclear architecture and Polycomb-dependent gene silencing. PLOS Genetics 16, e1008962 (2020).

43. T. Furuyama, R. Banerjee, T. R. Breen, P. J. Harte, SIR2 Is Required for Polycomb Silencing and Is Associated with an E(Z) Histone Methyltransferase Complex. Current Biology 14, 1812–1821 (2004).

44. A. Lagarou et al., dKDM2 couples histone H2A ubiquitylation to histone H3 demethylation during Polycomb group silencing. Genes Dev 22, 2799–2810 (2008).

45. R. P. Lifton, M. L. Goldberg, R. W. Karp, D. S. Hogness, The organization of the histone genes in Drosophila melanogaster: functional and evolutionary implications. Cold Spring Harb Symp Quant Biol 42 Pt 2, 1047–1051 (1978).

46. A. Chaouch et al., Histone H3.3 K27M and K36M mutations de-repress transposable elements through perturbation of antagonistic chromatin marks. Molecular Cell 81, 4876-4890.e4877 (2021).

47. C. Lu et al., Histone H3K36 mutations promote sarcomagenesis through altered histone methylation landscape. Science 352, 844–849 (2016).

48. R. Popovic et al., Histone methyltransferase MMSET/NSD2 alters EZH2 binding and reprograms the myeloma epigenome through global and focal changes in H3K36 and H3K27 methylation. PLoS Genet 10, e1004566 (2014).

49. P. J. Skene, J. G. Henikoff, S. Henikoff, Targeted in situ genome-wide profiling with high efficiency for low cell numbers. Nat Protoc 13, 1006–1019 (2018).

50. G. A. Orsi et al., High-resolution mapping defines the cooperative architecture of Polycomb response elements. Genome Research 24, 809–820 (2014).

51. J. G. Henikoff, J. A. Belsky, K. Krassovsky, D. M. Macalpine, S. Henikoff, Epigenome characterization at single base-pair resolution. Proceedings of the National Academy of Sciences 108, 18318–18323 (2011).

52. P. J. Skene, S. Henikoff, An efficient targeted nuclease strategy for high-resolution mapping of DNA binding sites. Elife 6, (2017).

53. N. Nègre et al., A cis-regulatory map of the Drosophila genome. Nature 471, 527–531 (2011).

54. J. Erceg et al., Dual functionality of cis-regulatory elements as developmental enhancers and Polycomb response elements. Genes Dev 31, 590–602 (2017).

55. B. A. Bredesen, M. Rehmsmeier, DNA sequence models of genome-wide Drosophila melanogaster Polycomb binding sites improve generalization to independent Polycomb Response Elements. Nucleic Acids Research 47, 7781–7797 (2019).

56. T. Fiedler, M. Rehmsmeier, jPREdictor: a versatile tool for the prediction of cis-regulatory elements. Nucleic Acids Res 34, W546–550 (2006).

57. L. Ringrose, M. Rehmsmeier, J.-M. Dura, R. Paro, Genome-Wide Prediction of Polycomb/Trithorax Response Elements in Drosophila melanogaster. Developmental Cell 5, 759–771 (2003).

58. J. Zeng, B. D. Kirk, Y. Gou, Q. Wang, J. Ma, Genome-wide polycomb target gene prediction in Drosophila melanogaster. Nucleic Acids Research 40, 5848–5863 (2012).

59. J. L. Brown, M.-A. Sun, J. A. Kassis, Global changes of H3K27me3 domains and Polycomb group protein distribution in the absence of recruiters Spps or Pho. Proceedings of the National Academy of Sciences 115, E1839–E1848 (2018).

60. M. I. Kuroda, H. Kang, S. De, J. A. Kassis, Dynamic Competition of Polycomb and Trithorax in Transcriptional Programming. Annual Review of Biochemistry 89, 235–253 (2020).

61. A. M. Deaton et al., Enhancer regions show high histone H3.3 turnover that changes during differentiation. eLife 5, (2016).

62. Y. Mito, J. G. Henikoff, S. Henikoff, Histone replacement marks the boundaries of cisregulatory domains. Science 315, 1408–1411 (2007).

63. A. Busturia et al., The MCP silencer of the<i>Drosophila Abd-B</i>gene requires both Pleiohomeotic and GAGA factor for the maintenance of repression. Development 128, 2163–2173 (2001).

64. B. A. Horard, C. Tatout, S. Poux, V. Pirrotta, Structure of a Polycomb Response Element and In Vitro Binding of Polycomb Group Complexes Containing GAGA Factor. Molecular and Cellular Biology 20, 3187–3197 (2000).

65. R. K. Mishra et al., The iab-7 Polycomb Response Element Maps to a Nucleosome-Free Region of Chromatin and Requires Both GAGA and Pleiohomeotic for Silencing Activity. Molecular and Cellular Biology 21, 1311–1318 (2001).

66. A. Schwendemann, M. Lehmann, Pipsqueak and GAGA factor act in concert as partners at homeotic and many other loci. Proceedings of the National Academy of Sciences 99, 12883–12888 (2002).

67. T. Nakayama, K. Nishioka, Y.-X. Dong, T. Shimojima, S. Hirose, <i>Drosophila</i> GAGA factor directs histone H3.3 replacement that prevents the heterochromatin spreading. Genes & Development 21, 552–561 (2007).

68. H. Okulski, B. Druck, S. Bhalerao, L. Ringrose, Quantitative analysis of polycomb response elements (PREs) at identical genomic locations distinguishes contributions of PRE sequence and genomic environment. Epigenetics & Chromatin 4, 4 (2011).

69. K. Ahmad, S. Henikoff, Histone H3 variants specify modes of chromatin assembly. Proc Natl Acad Sci U S A 99 Suppl 4, 16477–16484 (2002).

70. H.-T. Fang et al., Global H3.3 dynamic deposition defines its bimodal role in cell fate transition. Nature Communications 9, (2018).

71. E. McKittrick, P. R. Gafken, K. Ahmad, S. Henikoff, Histone H3.3 is enriched in covalent modifications associated with active chromatin. Proc Natl Acad Sci U S A 101, 1525–1530 (2004).

72. I. Bajusz et al., The Trithorax-mimic allele of Enhancer of zeste renders active domains of target genes accessible to polycomb-group-dependent silencing in Drosophila melanogaster. Genetics 159, 1135–1150 (2001).

73. C. T. Wu, M. Howe, A genetic analysis of the Suppressor 2 of zeste complex of Drosophila melanogaster. Genetics 140, 139–181 (1995).

74. I. A. Hernández-Romero, V. J. Valdes, De Novo Polycomb Recruitment and Repressive Domain Formation. Epigenomes 6, (2022).

75. R. K. Maeda, F. Karch, The ABC of the BX-C: the bithorax complex explained. Development 133, 1413–1422 (2006).

76. R. J. Diederich, V. K. Merrill, M. A. Pultz, T. C. Kaufman, Isolation, structure, and expression of labial, a homeotic gene of the Antennapedia Complex involved in Drosophila head development. Genes Dev 3, 399–414 (1989).

77. M. P. Scott et al., The molecular organization of the Antennapedia locus of Drosophila. Cell35, 763–776 (1983).

78. C. Ballare et al., Phf19 links methylated Lys36 of histone H3 to regulation of Polycomb activity. Nat Struct Mol Biol 19, 1257–1265 (2012).

79. L. Cai et al., An H3K36 methylation-engaging Tudor motif of polycomb-like proteins mediates PRC2 complex targeting. Mol Cell 49, 571–582 (2013).

80. C. A. Musselman et al., Molecular basis for H3K36me3 recognition by the Tudor domain of PHF1. Nat Struct Mol Biol 19, 1266–1272 (2012).

81. R. Cao et al., Role of hPHF1 in H3K27 Methylation and Hox Gene Silencing. Molecular and Cellular Biology 28, 1862–1872 (2008).

82. J. Choi et al., DNA binding by PHF1 prolongs PRC2 residence time on chromatin and thereby promotes H3K27 methylation. Nat Struct Mol Biol 24, 1039–1047 (2017).

83. K. Sarma, R. Margueron, A. Ivanov, V. Pirrotta, D. Reinberg, Ezh2 requires PHF1 to efficiently catalyze H3 lysine 27 trimethylation in vivo. Mol Cell Biol 28, 2718–2731 (2008).

84. A. Friberg, A. Oddone, T. Klymenko, J. Muller, M. Sattler, Structure of an atypical Tudor domain in the Drosophila Polycomblike protein. Protein Sci 19, 1906–1916 (2010).

85. S. O’Connell et al., Polycomblike PHD fingers mediate conserved interaction with enhancer of zeste protein. J Biol Chem 276, 43065–43073 (2001).

86. F. Tie, J. Prasad-Sinha, A. Birve, A. Rasmuson-Lestander, P. J. Harte, A 1-megadalton ESC/E(Z) complex from Drosophila that contains polycomblike and RPD3. Mol Cell Biol 23, 3352–3362 (2003).

87. M. Nekrasov et al., Pcl-PRC2 is needed to generate high levels of H3-K27 trimethylation at Polycomb target genes. The EMBO Journal 26, 4078–4088 (2007).

88. L. A. Banaszynski et al., Hira-dependent histone H3.3 deposition facilitates PRC2 recruitment at developmental loci in ES cells. Cell 155, 107–120 (2013).

89. B. Papp, J. Muller, Histone trimethylation and the maintenance of transcriptional ON and OFF states by trxG and PcG proteins. Genes Dev 20, 2041–2054 (2006).

90. J. van Arensbergen et al., A distal intergenic region controls pancreatic endocrine differentiation by acting as a transcriptional enhancer and as a polycomb response element. PLoS One 12, e0171508 (2017).

91. H. Lindehell, A. Glotov, E. Dorafshan, Y. B. Schwartz, J. Larsson, The role of H3K36 methylation and associated methyltransferases in chromosome-specific gene regulation. Sci Adv 7, eabh4390 (2021).

92. K. J. Evans et al., Stable Caenorhabditis elegans chromatin domains separate broadly expressed and developmentally regulated genes. Proc Natl Acad Sci U S A 113, E7020–E7029 (2016).

93. M. A. Willcockson et al., H1 histones control the epigenetic landscape by local chromatin compaction. Nature 589, 293–298 (2021).

94. F. Bantignies et al., Polycomb-dependent regulatory contacts between distant Hox loci in Drosophila. Cell 144, 214–226 (2011).

95. P. Buchenau, J. Hodgson, H. Strutt, D. J. Arndt-Jovin, The Distribution of Polycomb-Group Proteins During Cell Division and Development in Drosophila Embryos: Impact on Models for Silencing. Journal of Cell Biology 141, 469–481 (1998).

96. C. Grimaud et al., RNAi Components Are Required for Nuclear Clustering of Polycomb Group Response Elements. Cell 124, 957–971 (2006).

97. C. J. Lin, M. Conti, M. Ramalho-Santos, Histone variant H3.3 maintains a decondensed chromatin state essential for mouse preimplantation development. Development 140, 3624–3634 (2013).

98. Y. Wang et al., Histone variants H2A.Z and H3.3 coordinately regulate PRC2-dependent H3K27me3 deposition and gene expression regulation in mES cells. BMC Biology 16, (2018).

99. K. M. Dorighi, J. W. Tamkun, The trithorax group proteins Kismet and ASH1 promote H3K36 dimethylation to counteract Polycomb group repression in Drosophila. Development 140, 4182–4192 (2013).

100. Y. Tanaka, Z. Katagiri, K. Kawahashi, D. Kioussis, S. Kitajima, Trithorax-group protein ASH1 methylates histone H3 lysine 36. Gene 397, 161–168 (2007).

101. S. Schmahling et al., Regulation and function of H3K36 di-methylation by the trithorax-group protein complex AMC. Development 145, (2018).

102. T. Klymenko, J. Muller, The histone methyltransferases Trithorax and Ash1 prevent transcriptional silencing by Polycomb group proteins. EMBO Rep 5, 373–377 (2004).

103. U. Gunesdogan, H. Jackle, A. Herzig, A genetic system to assess in vivo the functions of histones and histone modifications in higher eukaryotes. EMBO Rep 11, 772–776 (2010).

104. S. Kondo, R. Ueda, Highly Improved Gene Targeting by Germline-Specific Cas9 Expression in Drosophila. Genetics 195, 715–721 (2013).

105. K. Ahmad, CUT&RUN with Drosophila tissues V.1. protocols.io. 2018 (dx.doi.org/10.17504/protocols.io.umfeu3n).

106. Y. Liao, G. K. Smyth, W. Shi, featureCounts: an efficient general purpose program for assigning sequence reads to genomic features. Bioinformatics 30, 923–930 (2014).

107. M. I. Love, W. Huber, S. Anders, Moderated estimation of fold change and dispersion for RNA-seq data with DESeq2. Genome Biology 15, (2014).

108. F. Ramírez et al., deepTools2: a next generation web server for deep-sequencing data analysis. Nucleic Acids Res 44, W160–165 (2016).

109. A. R. Quinlan, I. M. Hall, BEDTools: a flexible suite of utilities for comparing genomic features. Bioinformatics 26, 841–842 (2010).

110. S. Andrews. (Babraham Bioinformatics, Babraham Institute, Cambridge, United Kingdom, 2010).

111. S. W. Wingett, S. Andrews, FastQ Screen: A tool for multi-genome mapping and quality control. F1000Research 7, 1338 (2018).

112. B. Langmead, S. L. Salzberg, Fast gapped-read alignment with Bowtie 2. Nat Methods 9, 357–359 (2012).

113. P. Danecek et al., Twelve years of SAMtools and BCFtools. Gigascience 10, (2021).

114. W. J. Kent, A. S. Zweig, G. Barber, A. S. Hinrichs, D. Karolchik, BigWig and BigBed: enabling browsing of large distributed datasets. Bioinformatics 26, 2204–2207 (2010).

115. Y. Zhang et al., Model-based Analysis of ChIP-Seq (MACS). Genome Biology 9, R137 (2008).

116. M. N. Patwardhan, C. D. Wenger, E. S. Davis, D. H. Phanstiel, Bedtoolsr: An R package for genomic data analysis and manipulation. J Open Source Softw 4, (2019).

